# CRISPR comparison toolkit (CCTK): Rapid identification, visualization, and analysis of CRISPR array diversity

**DOI:** 10.1101/2022.07.31.502198

**Authors:** Alan J. Collins, Rachel J. Whitaker

## Abstract

CRISPR-Cas systems provide immunity against mobile genetic elements (MGEs) through sequence-specific targeting by spacer sequences encoded in CRISPR arrays. Spacers are highly variable between microbial strains and can be acquired rapidly, making them well suited for use in strain typing of closely related organisms. However, the historical record of spacer acquisitions and deletions represented by arrays has not previously been used to reconstruct strain histories. We therefore developed the CRISPR Comparison Toolkit (CCTK) to enable analyses of array relationships. CCTK includes tools to identify arrays, analyze relationships between arrays using CRISPRdiff and CRISPRtree, and predict targets of spacers. CRISPRdiff visualizes arrays and highlights the similarities between them. CRISPRtree infers a phylogenetic tree from array relationships and presents a hypothesis of the evolutionary history of the arrays. CCTK unifies several CRISPR analysis tools into a single command line application, including the first tool to infer phylogenies from array relationships.

## Introduction

CRISPR-Cas (clustered regularly interspersed short palindromic repeats; CRISPR-associated proteins) is an adaptive immune system present in most archaea and many bacteria that provides immunity against mobile genetic elements (MGEs) such as viruses ^1,2^. CRISPR-Cas immunity is acquired through the incorporation of spacers (small fragments of DNA from invading MGEs) into a region called a CRISPR array. CRISPR arrays are dynamic as they can both acquire and lose spacers. Over time, these events alter the length and spacer content of arrays ^3^. Spacer acquisition typically occurs at the leader end ^2^. However, under some circumstances, ectopic spacer acquisition can occur ^4,5^. Repeats located at the leader-distal end of the array (the trailer end) are thought to be older and sometimes contain polymorphisms ^6^. Because proper CRISPR function requires the recognition of specific sequence and secondary structure in the repeats ^7^, mutations within trailer-end repeats can result in the loss of function of spacers flanked by the degraded repeats. Spacers can also be lost by deletion events ^8,9^. Specifically, homologous recombination between repeat sequences can lead to the loss of a short stretch of spacers and repeats ^3,10,11^.

Because thousands of unique spacers can be derived from a single virus ^12,13^, there is an enormous number of possible spacer sequences. The presence of two identical spacers in different CRISPR arrays is unlikely to have arisen by chance and may indicate that the arrays share a common ancestor (i.e., the arrays are homologous). Therefore, comparisons of spacers between CRISPR arrays can reveal phylogenetic relationships between them.

To infer phylogenetic relationships between CRISPR arrays, one must first visualize and compare the arrays to one another. The previously published tools CRISPRviz and CRISPRStudio show which spacers are shared or distinct between CRISPR arrays ^14,15^; however, no tools have been developed to infer phylogenetic relationships between arrays.

Here, we present the CRISPR comparison toolkit (CCTK), a command-line software toolkit for the analysis of relationships between CRISPR arrays. CCTK includes tools to identify CRISPR arrays and predict protospacers. CCTK also includes two new programs for the analysis of array relationships: CRISPRdiff, which visualizes arrays and highlights regions of similarity, and CRISPRtree, which infers the phylogenetic relationships between a set of arrays using a maximum parsimony approach. By unifying the above tools into a single, command-line application, CCTK provides resources for identifying relationships between CRISPR arrays in a few simple steps. Furthermore, CCTK uses simple file formats to facilitate the integration of CCTK tools into existing pipelines.

## Methods

### Dataset description

We used CCTK to analyze CRISPR arrays encoded by *Pseudomonas aeruginosa*, and we present the results of those analyses in the following sections. Specifically, we analyzed a dataset that includes information about the phylogenetic relationships between *P. aeruginosa* isolates and the CRISPR arrays they encode ^16,17^. The flow of information between CCTK tools is shown in **Figure 1**. The sequence records analyzed here and the CRISPR arrays identified are described in **Table S1**.

**Figure 1.**
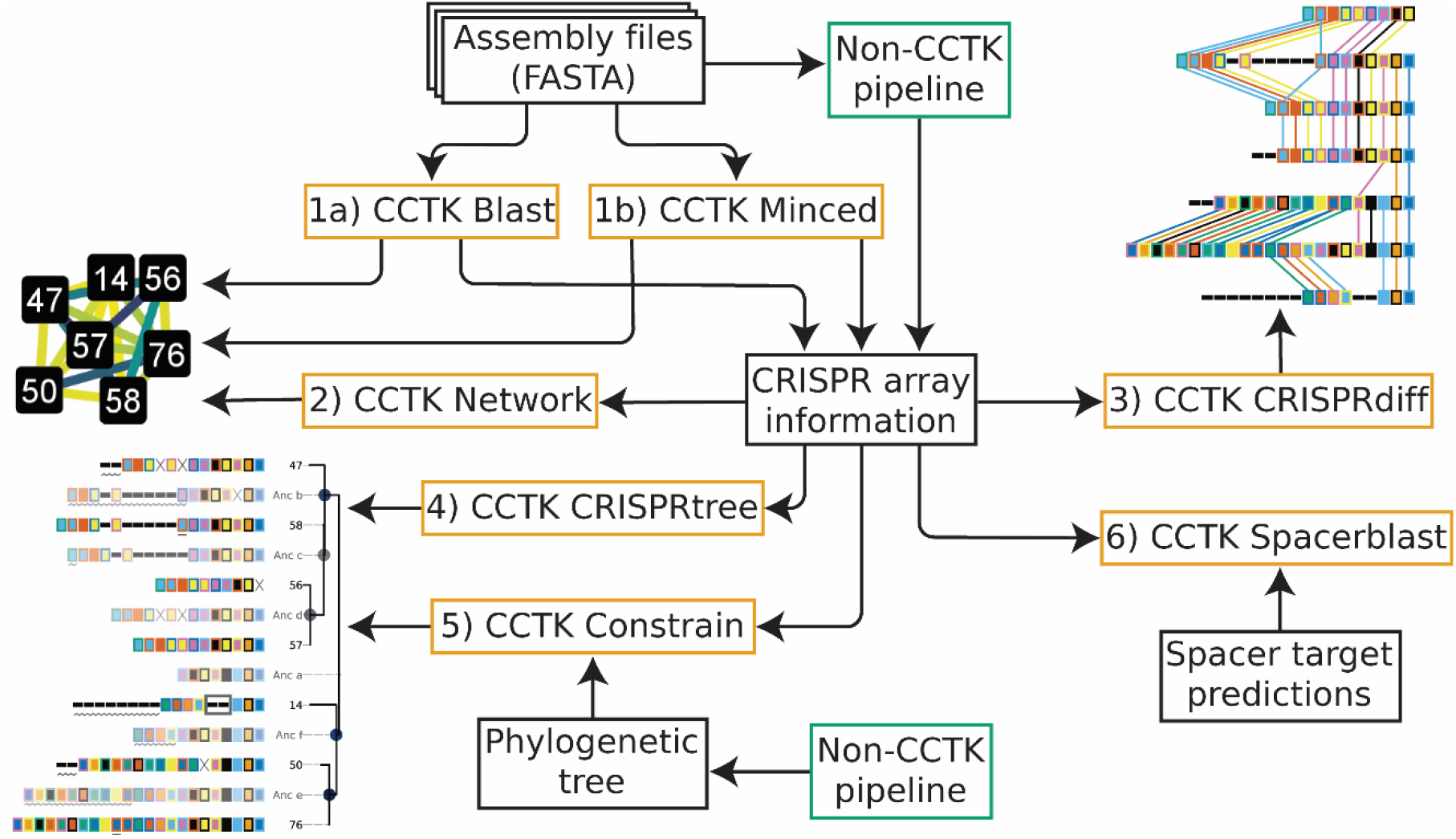
Flow of data through CCTK tools. Beginning with genome assemblies in FASTA format, CCTK tools can be used to identify and analyze CRISPR arrays. The CCTK tools are: 1a) Blast, 1b) Minced, 2) Network, 3) CRISPRdiff, 4) CRISPRtree, 5) Constrain, and 6) Spacerblast. The functions of each CCTK tool are as follows: 1) CRISPR arrays can be identified in assemblies using either MinCED or BLASTN followed by processing steps that produce aggregated CRISPR information for all input assemblies. These CRISPR information files are used by downstream CCTK tools. Alternatively, non-CCTK tools can be used to identify arrays, and their output easily adapted to the simple file formats used by CCTK. 2) The relationships between CRISPR arrays can be represented as a network in which each array is represented as a node. Homologous arrays (i.e., arrays containing identical spacers) are connected by an edge, the weight of which corresponds to the Jaccard similarity between the two arrays. 3) Groups of homologous arrays can be visualized using CRISPRdiff to identify the differences and similarities between arrays. 4) CRISPRtree can infer a phylogenetic tree representing the relationships between homologous arrays and can hypothesize events that occurred during their history. 5) Constrain can visually represent how CRISPR arrays may have evolved given a fixed tree topology to allow the reconciliation of CRISPR relationships with other phylogenetic data. 6) Spacerblast predicts protospacer targets of CRISPR spacers and determines the presence of a protospacer adjacent motif.

### Sequence assembly and core genome identification

The sequence reads associated with the 72 “clone-corrected” isolates described by England, et al., first published by Marvig et al., were retrieved from the European Nucleotide Archive ^16,17^. Assemblies were produced using Spades with the “careful” option ^18^. An alignment of the core genome sequences of these assemblies was identified using Spine, Nucmer, and a custom script ^19,20^(Supplemental Methods). IQTREE2 was used to infer a maximum likelihood tree from this core genome alignment (**Fig. S1**) ^21^.

### CRISPR array identification

CCTK includes two tools to identify CRISPR arrays in genome assemblies, CCTK Minced and CCTK Blast. CCTK Minced uses MinCED, which identifies CRISPR arrays using a sliding window search to identify regularly spaced repeats and can identify CRISPR arrays without requiring the user to have prior knowledge of the expected CRISPR subtypes ^22^. CCTK Blast uses BLASTN to identify CRISPR arrays by searching for a user-defined set of CRISPR repeat sequences ^23^. Both tools can use repeat orientation information (provided by the user) to correctly orient the identified arrays. Optionally, both tools can assess the sequence similarity between spacers to collapse groups of spacers into a single representative (when they differ by fewer than a threshold of bases specified by the user). Alternatively, CRISPR arrays identified using other approaches can be converted into the simple file format used by CCTK tools. Additional details about CCTK Blast and CCTK Minced are provided in the Supplemental Methods.

### CCTK Network representation of homologous CRISPR arrays

CCTK produces a network in which CRISPR arrays are represented as nodes (**Fig. 2**). Two arrays are connected by an edge if they contain identical spacers (i.e., are homologous). CCTK quantifies homology between two arrays using Jaccard similarity: the number of spacers in common between two arrays divided by the total number of unique spacers present in the two arrays ^24^.

**Figure 2.**
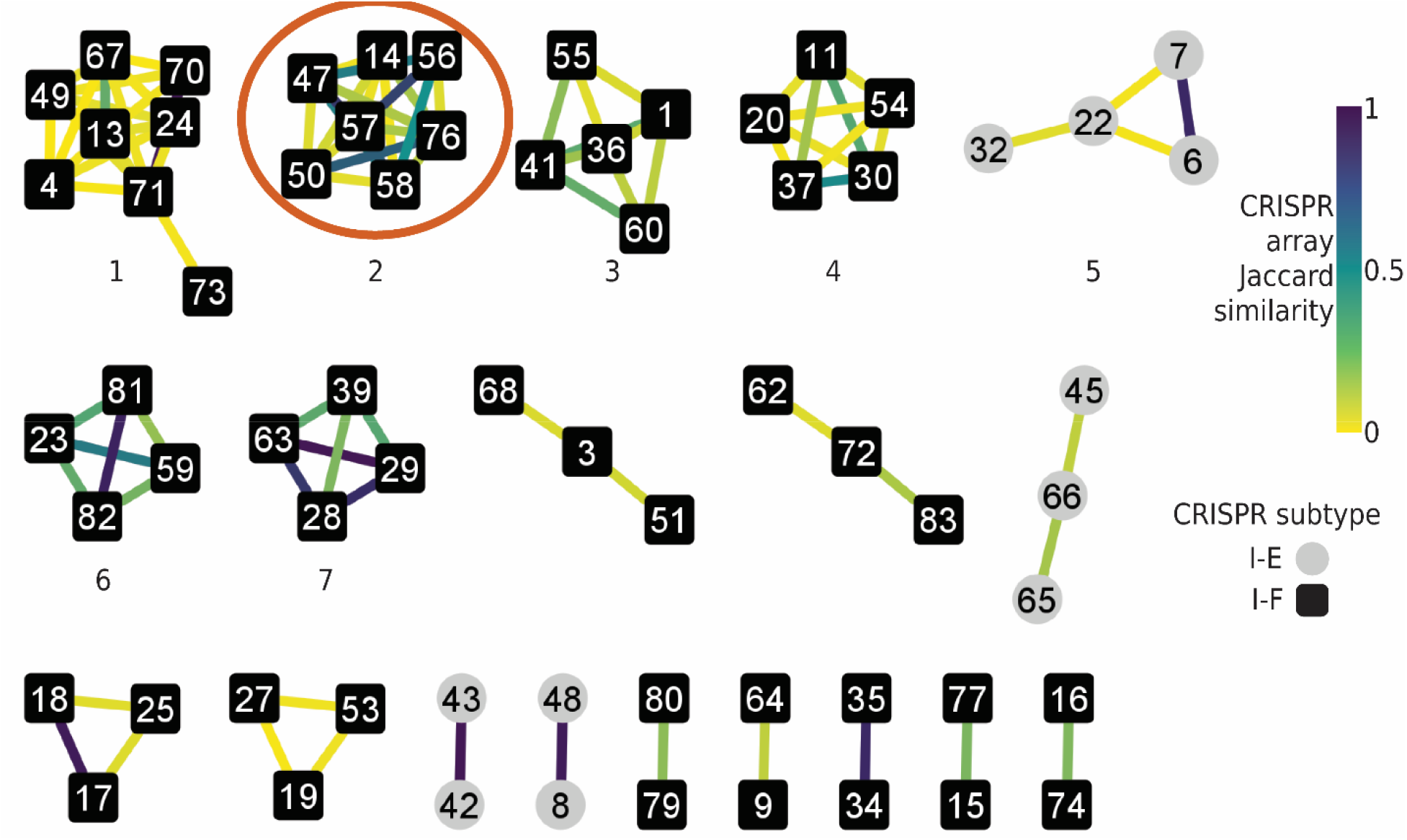
Clusters of homologous arrays can be identified using a network representation of array relationships. Network representation of spacers shared between arrays in *Pseudomonas aeruginosa* isolates clone corrected by England et al. ^16^, visualized using Cytoscape ^41^. Each CRISPR array is represented by a node. An edge is drawn between two nodes when CRISPR arrays share at least two spacers. Edge colour corresponds to the Jaccard similarity between the two connected arrays (i.e., the number of spacers shared between the two arrays divided by the number of total unique spacers in the two arrays) as indicated in the colour key. The seven largest clusters are numbered (corresponding to the cluster numbers in **Fig. S1**). Circled is a cluster of arrays, hereafter called cluster_2, which are analysed further in subsequent figures.

CCTK Minced and CCTK Blast each produce a network when they are used to identify arrays. Alternatively, Network produces a network representation of CRISPR arrays identified using any method (and provided in the format required by CCTK) (**Fig. 1**). For all three of these tools, the user can specify the minimum number of spacers that must be shared between two arrays for an edge to be drawn between them.

### CRISPRdiff: CRISPR array visualization

CRISPRdiff visualizes CRISPR arrays through three steps. First, spacers that are present in more than one array assigned a unique combination of fill and outline colour. Unique spacers are represented as thin, black rectangles (**Fig. 3**). Next, the order of arrays in the plot is determined to maximize the number of spacers shared between adjacently plotted arrays. Finally, the plot is drawn using Matplotlib, which allows the plot to be saved as many common file formats ^25^.

**Figure 3.**
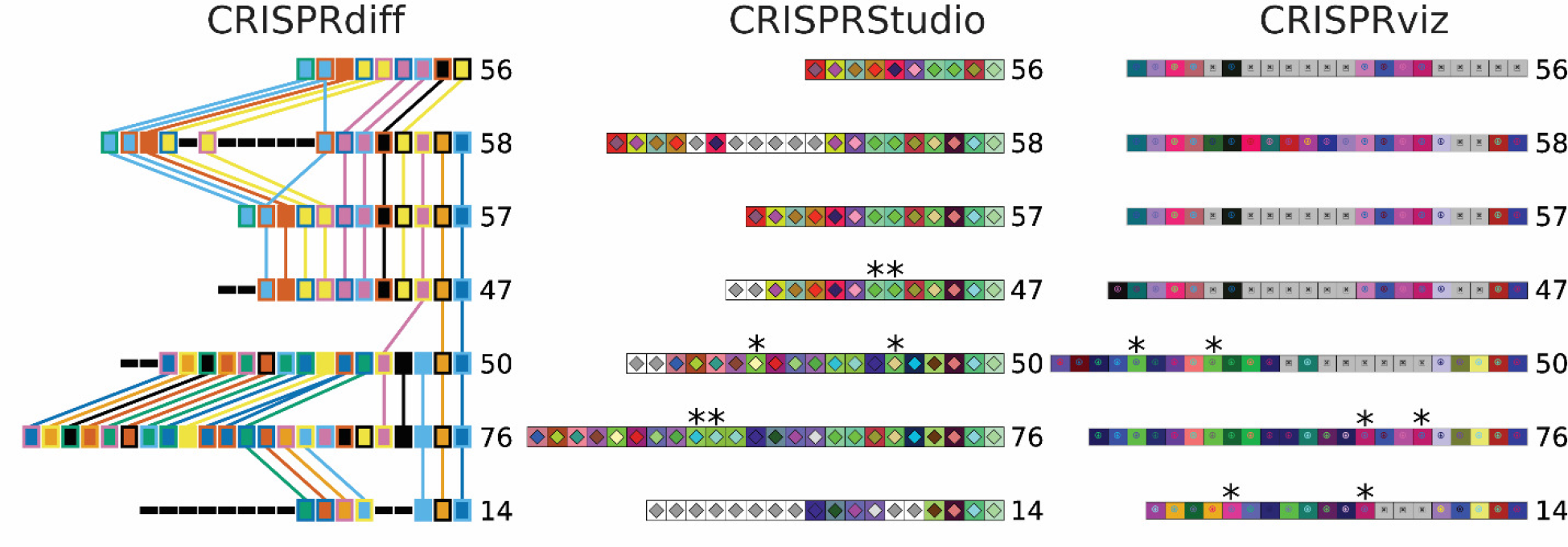
CRISPRdiff produces a clearer illustration of array relationships than previously published tools. Cluster_2 arrays were visualized using each of the following tools: CRISPRdiff, CRISPRStudio, and CRISPRviz. The leader end of each array is on the left and the trailer end is on the right. CRISPRdiff and CRISPRviz show arrays identified using MinCED ^22^, while CRISPRStudio shows arrays identified using its companion tool, CRISPRdetect ^42^. The plot produced by CRISPRStudio shows an additional spacer with ten mismatches at the trailer end of each array that was only identified by CRISPRdetect. CRISPRStudio was run using the –gU option to assign unique spacers a grey fill colour. The automatic spacer alignment function of CRISPRviz was used, followed by manual correction (gaps in the alignment are represented as grey boxes). The order of arrays in the images produced by CRISPRStudio and CRISPRviz was manually altered to correspond to the order chosen by CRISPRdiff. Array identifiers assigned by CCTK Minced are shown next to each corresponding array. “*” indicates spacers that were assigned colours with low visual contrast by CRISPRStudio or CRISPRviz.

Spacers are assigned colours using either a default colourblind-friendly colour scheme ^26^, or user-specified colours. Array order is determined by an approach that maximizes the number of spacers in common between adjacently plotted arrays. For small numbers of arrays, the best possible order is found. For larger numbers of arrays, a limited search is performed, and the best order found is used. The user can also set the array order. See the Supplemental Methods for additional details about colour assignment and array order determination.

### CRISPRtree: Maximum parsimony analysis of CRISPR array relationships

Given a list of arrays to be analyzed, CRISPRtree constructs a tree starting with two arrays and then adds the remaining arrays to the tree one by one. The process used by CRISPRtree (illustrated in **Fig. S2**) is as follows. First, two CRISPR arrays are aligned using the Needleman-Wunsch algorithm (**Fig. S2B**). Each spacer is treated as a character and aligned using the following scores: match: 100, mismatch: −1, gap: −2 ^27^. These scores were empirically chosen to ensure that shared spacers are always aligned. The aligned arrays are then split into modules. Modules are classified by identifying sets of consecutive spacers that have the same relationship between the aligned arrays (**Fig. S2B**). For example, a set of consecutive spacers present at the leader end of one array, but missing in the other, is classified as a “leader-end acquisition module.” A set of consecutive spacers present in both arrays is classified as a “shared module.”

Next, a hypothetical ancestor array is generated by using an evolutionary model of CRISPR array change that involves the following possible events: leader-end acquisition, the loss of spacers one at a time from the trailer end, and deletion or insertion of one or more spacers anywhere in the array (**Fig. S2C**). The process by which CRISPRtree processes an array alignment to infer an ancestral array is described in the Supplemental Methods.

After a hypothetical ancestral array has been inferred, events in each descendant array are identified and parsimony costs assigned. Events are identified by processing each descendant of the newly generated ancestral array as follows (**Fig. S2B)**. First, the descendant array and ancestor array are aligned. Then, modules of spacers are identified in the descendant array as described above (**Fig. S2B**). Next, each module is associated with a type of event (i.e., acquisition, insertion, trailer loss, etc.) Finally, the parsimony costs for all identified events are summed. The total parsimony cost of events identified in a descendant array are set as the branch length between the array and its ancestor in the tree. Default event parsimony costs were empirically determined. Default event parsimony costs are acquisition: 1, duplication: 1, insertion: 30, deletion: 10, trailer loss: 1, independent acquisition: 50. These costs can also be set by the user.

Once the tree has been initialized with the first two arrays and their hypothetical ancestral array (**Fig. S2D**), additional arrays are added to the tree. To add an array to the tree, it is first compared to each array already present in the tree (**Fig. S2E**). The closest match already present in the tree is then set as the sibling node of the newly added array and a hypothetical ancestor is generated (**Fig. S2F**,**G**,**H**).

When all arrays have been added to the tree, any ancestral arrays that are themselves identical to their ancestor (i.e., internal nodes with branch length of zero) are collapsed to a polytomy. Tree manipulation is performed using the python package DendroPy ^28^.

Different trees are generated in this manner by adding arrays to the tree in different orders. Each tree is then scored according to the total branch length of the tree (i.e., the total parsimony cost of all hypothesized events). The tree (or trees if multiple equally parsimonious trees are found) with the lowest total parsimony cost is considered the best. Trees are output in Newick format and visualized using Matplotlib ^25^.

Optionally, a measure of node support can be calculated by CRISPRtree. Node support is calculated using the replicates of the tree building process used by CRISPRtree. First, each input array is assigned a position in a binary string. Then, every tree produced by CRISPRtree is encoded as a list of binary strings. Each element in the list represents an internal node in the tree and the binary string indicates which arrays are descendants of the node. For example, for three arrays A, B, and C (which are assigned positions in a binary string in alphabetical order), an internal node with A and C as descendants would be encoded as 101. Once all trees are encoded in this way, the nodes in the most parsimonious tree(s) are compared to all other trees, and the number of times that each node is seen is counted. This count is then divided by the total number of trees constructed, resulting in a proportion of trees containing that node. Node support is reported in the Newick string and indicated using a coloured circle at the corresponding internal node in the graphical representation of the tree.

### Constrain: Inference of CRISPR array evolution given a fixed tree topology

Given a phylogenetic tree, Constrain infers an evolutionary history for a set of arrays that fits the provided topology. The process used by Constrain is as follows. First, the leaves on the input tree are populated with CRISPR arrays. Then, starting with leaf nodes and working towards the root of the tree, the array at each node is aligned with the array at its sibling node and an ancestral array is hypothesized using the same approach as that used by CRISPRtree (illustrated in **Fig. S2B**,**C**). Once all internal nodes in the tree have been populated with hypothetical ancestral arrays, each array is annotated with events that would have occurred for it to have arisen from its ancestral array.

### Tree comparisons

The arrays produced by each simulation performed using Evolve (See Supplemental Methods) were analyzed with CRISPRtree. The CRISPRtree output was compared to the true tree (produced by Evolve) using Robinson-Foulds (RF) distance, which measures the number of changes it would take to convert one tree into another ^29^. RF distance was calculated using the ete3 toolkit version 3.1.2 ^30^ in python3.8.12 ^31^.

To compare the RF distance between simulation replicates (in which trees with different numbers of nodes may be produced), each distance is expressed as a fraction of the maximum possible RF distance between the two trees, indicating the extent to which the two trees differ relative to two completely distinct trees.

This analysis was performed for simulations run with different parameters: numbers of events (e.g., the acquisition of a spacer), which controls the size and extent of divergence between arrays; the rate of loss of arrays, which impacts the level of incompleteness of the dataset and the extent to which extant arrays are diverged; and the relative rates of spacer acquisition and deletion, which determines the type of relationships between arrays and, at high levels of deletion, the extent to which the relationships between arrays is obscured by spacer loss. Parameters used in these simulations are described in more detail in the Supplemental Methods section.

### Software availability

All source code is available on GitHub https://github.com/Alan-Collins/CRISPR_comparison_toolkit. CCTK can be installed either by downloading from the GitHub repository or using the package manager Anaconda https://anaconda.org/bioconda/cctk. Documentation describing usage of CCTK tools is available at https://crispr-comparison-toolkit.readthedocs.io/en/latest/.

CCTK relies on the following dependencies (version numbers in use at publication): Python3.8, Dendropy 4.5.2, Matplotlib 3.5.0, NumPy 1.21.2, MinCED 0.4.2, and BLAST+ 2.12.0+ ^22,23,31–33^.

## Results

### CRISPRdiff compares arrays and visualizes the relationships between them

In this study, we analyzed a dataset of previously described *P. aeruginosa* isolates ^16,17^. The phylogenetic relationships between these isolates are shown in **Figure S1**. In the following sections, we present analyses using CCTK to characterize the relationships between CRISPR arrays encoded by these *P. aeruginosa* isolates.

We first used CCTK Minced to identify CRISPR arrays (**Table S1**) and construct a network representing the sharing of spacers between homologous arrays (**Fig. 2**). The cluster of arrays circled in **Figure 2** (referred to as cluster_2 arrays) will be used to illustrate the functionality of CCTK tools. These arrays correspond to “RAG 3” described by England et al. ^16^. Except where stated, all subsequent analyses presented will focus on cluster_2 arrays.

Visualizing array relationships as a network enables the easy recognition of sets of homologous and unrelated arrays. Comparison of the spacers that are shared between homologous arrays can then be used to further explore the relationships between those arrays. There are two previously developed tools for visualization and comparison of CRISPR arrays: CRISPRviz and CRISPRstudio ^14,15^. While both tools can be used to produce visual representations of CRISPR arrays, they have limitations when used to identify similarities among a group of arrays. Therefore, we developed a new visualization tool, CRISPRdiff, as part of CCTK, to more effectively identify the differences and similarities between arrays.

Key elements that differ between CRISPRdiff, CRISPRstudio, and CRISPRviz are described in **Table S2**, and the visualizations produced by the three tools are shown in **Figure 3**. The most significant advantages of CRISPRdiff over CRISPRstudio and CRISPRviz are discussed further here.

The ability to identify shared spacers is important when comparing homologous arrays. CRISPRdiff uses lines connecting identical spacers to clearly highlight the presence of shared spacers. In addition, CRISPRdiff uses a colourblind-friendly colour palette that ensures visually distinct colours are assigned to different spacers ^26^. Alternatively, CRISPRdiff allows the user to define their own colour scheme. CRISPRStudio and CRISPRviz produce visualizations that may include colours that are not visually distinct or colourblind-friendly (spacers indicated by asterisks, **Fig. 3; Fig. S2; Table S2**), which makes it difficult to identify shared spacers.

Unlike the other tools, no further processing of the output image is required for CRISPRdiff. Instead, which arrays are included in the plot produced by CRISPRdiff can be specified in the command line. In contrast, when using CRISPRStudio or CRISPRviz, the user must manually process the output images to remove unwanted arrays that were identified in the input assemblies. Furthermore, in the case of CRISPRviz the user is also required to manually experiment with reverse-complement and reversing the order of spacers in arrays to visually identify homologous arrays.

### CRISPRtree infers array relationships using maximum parsimony

CRISPR arrays preserve a record of the MGEs encountered by a cell. In addition, the changes that have occurred within a CRISPR array over time represent a history of the evolution of that array. Comparing homologous CRISPR arrays and reconstructing the events that may have occurred during their evolution allows us to infer the phylogenetic relationships between CRISPR arrays and the genomes in which they are located. However, no tools have previously been developed to analyze the phylogenetic relationships between homologous CRISPR arrays. We therefore developed CRISPRtree, which infers the phylogenetic relationships among a set of arrays using a maximum parsimony approach (see Methods and Supplemental Methods; **Fig. S2**).

**Figure 4A** shows the tree inferred by CRISPRtree for cluster_2 arrays. CRISPRtree hypothesizes that all seven extant arrays descended from a last common ancestor, Anc_a, containing 8 spacers. Using the events annotated on each CRISPR array in **Figure 4A**, the hypothetical history of the analyzed arrays produced by CRISPRtree can be reconstructed.

**Figure 4.**
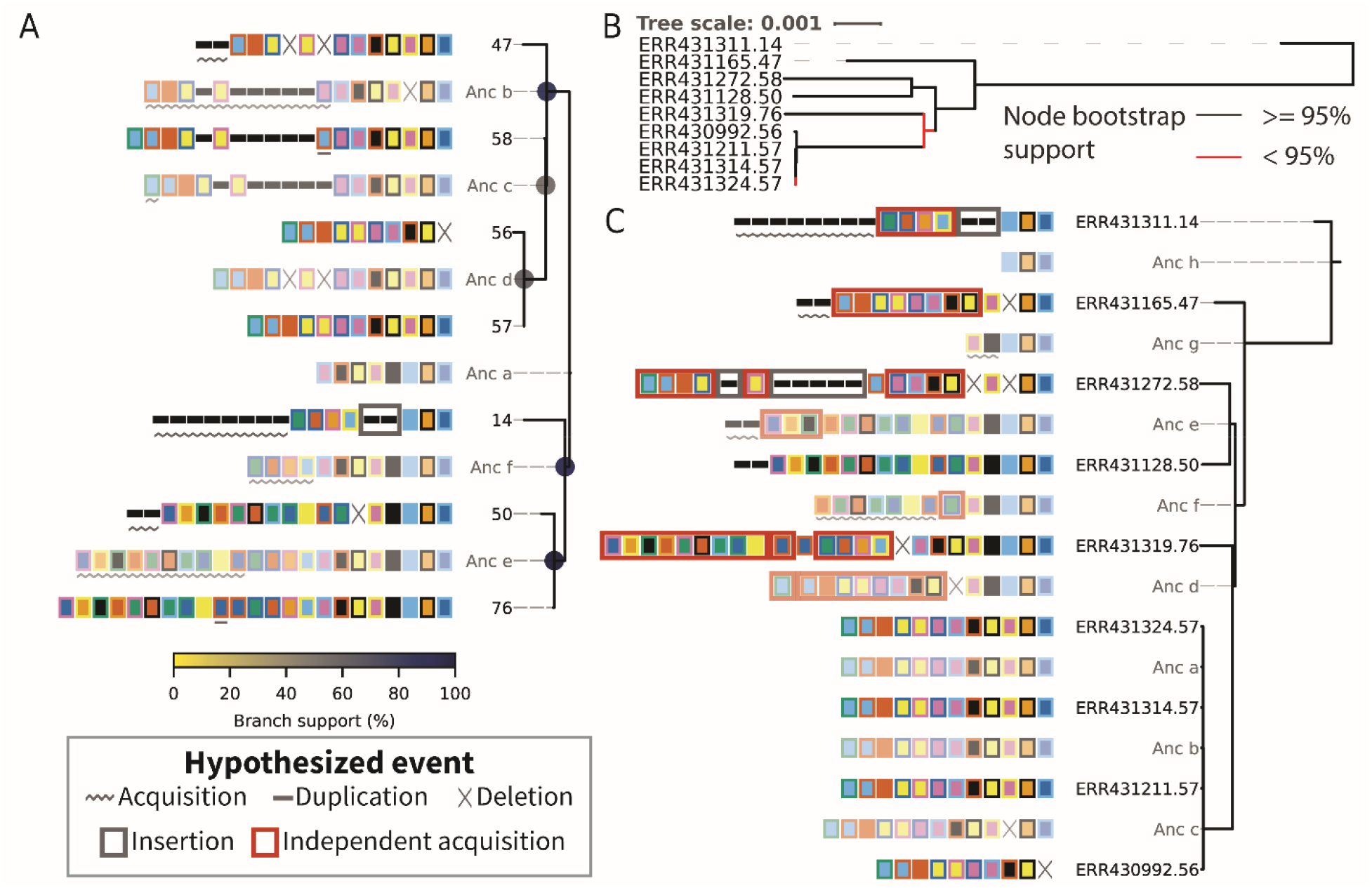
CRISPRtree can infer a phylogenetic tree from array relationships and Constrain can assess other phylogenetic data. (**A**) CRISPRtree was used to infer a tree representing the relationships between cluster_2 arrays. Leaf nodes in the tree correspond to the seven extant arrays (i.e., the arrays provided as input to CRISPRtree). Each internal node in the tree corresponds to a hypothetical ancestral state of the descendent arrays. Hypothetical ancestors are assigned names beginning with “Anc” and are rendered partially transparent to provide visual contrast. Next to each node label is a visual representation of the corresponding CRISPR array. Visualizations are annotated with events that are hypothesized to have occurred since the ancestor of that array (see “Hypothesized event” box). No events are highlighted on the root node array (Anc_a) as it is the last common ancestor of all extant arrays being considered and does not have an ancestral state. Node support is indicated using coloured circles at the corresponding branch (see colour scale below tree) (**B**) Maximum likelihood tree for the isolates encoding Cluster_2 arrays. Core genome SNPs were identified using Spine, Nucmer, and a custom script (Supplemental Methods) ^19,20^. IQTREE2 was then used to infer a tree using a model selected by ModelFinder (UNREST+FO+R3) ^21,43^. Red branches indicate node support below 95% calculated using the Ultrafast bootstrap tool included in IQTREE2 ^37^. Leaf names are composed of the European Nucleotide Archive accession number and the cluster_2 array ID encoded by that isolate. (**C**) Tree produced by Constrain representing the events that Constrain hypothesized would have to have occurred if cluster_2 arrays had evolved according to the core genome phylogeny shown in panel **B**. Events are highlighted using symbols described in the key. Leaf labels correspond to those in panel **B** and indicate the assembly accession and cluster_2 array ID.

From Anc_a, two arrays arose: Anc_b and Anc_f. The events in the upper clade, starting with Anc_b, will be described here to illustrate how the visualization produced by CRISPRtree can be interpreted. Anc_b differs from its ancestor Anc_a by the deletion of two spacers and the acquisition of several. From Anc_b, the extant array 47 arose by the deletion of two sets of spacers and the acquisition of two spacers, and Anc_c arose through the acquisition of a single spacer. From Anc_c, the extant array 58 arose through the duplication of an existing spacer, and Anc_d arose through two deletions. The extant array 57 has no annotated events, indicating that it is identical to Anc_d, while array 56 differs from Anc_d by a single deletion. As array 57 and Anc_d are identical, array 57 can be considered to be the ancestor of array 56.

CRISPRtree uses an evolutionary model that favours spacer gain by leader-end acquisitions, and spacer loss by deletions or trailer-end loss. However, sometimes other events can be predicted. For example, in the lower clade, Array 14 is hypothesized to have replaced five spacers in its middle with two new spacers. While this is annotated as an insertion event, it is also possible that the two spacers annotated as an insertion here may instead have been present in Anc_f and Anc_a, but independently lost in both Anc_b and Anc_e. These two mutually exclusive explanations for how the set of extant arrays may have arisen highlights the importance of being able to see and assess the evidence used to generate a tree topology. By clearly showing how it arrived at a given topology, CRISPRtree makes it easy for a user to decide whether the tree is a reasonable hypothesis or not.

In addition to illustrating the ancestral arrays and events, CRISPRtree can assign a measure of node support to each internal node in the tree (described in the Methods section). This node support value indicates the frequency with which a given node was seen in the trees generated during CRISPRtree’s search for the most parsimonious tree. For example, the node support at Anc_e is 100%, indicating that arrays 50 and 76 were placed together in every tree inferred by CRISPRtree during its search. The node support at Anc_c however, is 52%, indicating that in almost half of the trees inferred by CRISPRtree, a node containing only arrays 56, 57, and 58 was not present. These node support values provide an indication of how strong the phylogenetic signal is that places a set of arrays together given the evolutionary model used by CRISPRtree.

### Constrain can identify signals of horizontal gene transfer of CRISPR arrays

A common approach to examine the possibility of horizontal gene transfer (HGT) is phylogenetic reconciliation. Phylogenetic reconciliation involves the comparison of the tree of an individual gene’s evolution to a species tree ^34^. If the topology of a gene tree differs from the species tree, that is evidence that the gene has been horizontally transferred.

When comparing trees, it is important to assess both whether the two trees are different, and whether each tree could also explain the data underlying the other. Numerous statistical tests have been developed to test whether different tree topologies could explain the same nucleotide sequence alignment data. For example, the Shimodaira-Hasegawa (SH) test is commonly used to assess whether a set of trees are all good explanations of a sequence alignment ^35,36^.

We developed Constrain to assess whether different tree topologies may be good explanations of the array relationships. Constrain reconstructs the history of a set of CRISPR arrays given a certain tree topology and indicates the events that would need to occur during the evolution of each array. While Constrain is not a statistical testing method, it can indicate whether the given topology is reasonable or whether HGT or the independent acquisition of identical spacers in different arrays would be the most parsimonious explanation of the tree topology.

To investigate whether Cluster_2 arrays may have been horizontally transferred between isolates, we first determined whether the CRISPR array tree and core genome tree differ (**Fig. S1, 4A, B**). We used Robinson-Foulds (RF) distance (which assesses the number of operations required to convert one tree into another) to compare the tree produced by CRISPRtree to the core genome tree ^29^. The RF distance between the two trees is 9 out of a maximum possible value of 13. The high RF distance between the two trees indicates that there is a high degree of disagreement between the inferred histories of Cluster_2 CRISPR arrays and the core genome of the isolates encoding these arrays. Disagreement between the two trees is consistent with the hypothesis that horizontal gene transfer of CRISPR arrays has occurred.

We next assessed whether differences between the two trees are well supported by the underlying data. Bootstrap support for two of the nodes in the core genome tree is low (bootstrap support <95% - a cutoff recommended for use with UFBoot2 values ^37^). In addition, two nodes in the CRISPRtree have low support (Anc_c: 52%, Anc_d: 60%). However, major differences between the two trees are well supported. For example, Anc_f in the tree inferred by CRISPRtree places arrays 14, 50, and 76 together with 88% support. The core genome tree places the isolates encoding those three arrays (ERR431311.14, ERR431128.50, and ERR431319.76 in **Fig. 4B**) apart from one another with high bootstrap support. The finding that major differences between the two trees are well supported by the underlying data provides confidence that the two trees are different.

We next determined whether the topology of the CRISPRtree tree is supported by the core genome sequence alignment using the SH test implemented in IQTREE2 ^21,35^. The SH test indicates that the CRISPRtree topology for cluster_2 arrays is not supported by the alignment of core genomes (P=0). This indicates that the CRISPR array tree does not represent the relationships between the core genome sequences of these isolates.

Finally, we used Constrain to evaluate whether the core genome tree could explain the relationships between Cluster_2 CRISPR arrays. Constrain indicates that several spacers would need to have been independently acquired multiple times (**Fig. 4C**). These independent acquisition events are indicated as a red box surrounding the sets of spacers that Constrain identified as also being acquired in a different array. Independent acquisition events are annotated whenever one or more spacers could have been horizontally transferred and added to an array by recombination or independently added to multiple arrays through the normal CRISPR adaptation process.

Constrain indicates that the core genome tree does not explain the relationships between Cluster_2 arrays. Additionally, the distribution of cluster_2 CRISPR arrays among assemblies in this dataset provides evidence of HGT (**Fig. S1**). Specifically, Isolates encoding cluster_2 arrays are distantly related and, in some cases, more closely related to isolates that either do not encode cluster_2 arrays or have no detectable CRISPR arrays. Taken together, these observations are consistent with a hypothesis that cluster_2 arrays have been horizontally transferred between divergent isolates.

### Identifying spacer targets using Spacerblast

As cluster_2 arrays appear to have been involved in HGT events, and these isolates were sampled from a single patient population ^17^, we reasoned that arrays may spread by HGT as they provide immunity against phages or other MGEs that are present.

CRISPR systems provide immunity against MGEs through sequence-specific targeting ^2^. In addition, CRISPR targeting often requires the presence of a short sequence flanking the target called the protospacer adjacent motif (PAM) ^38,39^. The identification of CRISPR spacer targets therefore involves searching for sequence with high identity to the spacer that is flanked by a PAM. We developed a tool as part of CCTK, called Spacerblast, that can search for spacer matches in a sequence database and identify the presence of PAMs (Supplemental Methods). We next used Spacerblast to assess whether the spacers in cluster_2 arrays have targets within the assemblies in our dataset.

We searched for spacer matches with 100% sequence identity and the presence of a type I-F “CC” PAM on the 5’ flank of the protospacer ^39^. Spacerblast identified 77 predicted protospacers with flanking PAMs (**Table S3**). Each array in cluster_2 has at least one spacer with several predicted protospacers in the assemblies in this dataset (**Fig. 5A**). In addition, the cluster_2 array spacers with the most targets are predicted by Constrain to have been independently acquired by multiple arrays (**Fig. 5B**). These data are consistent with the hypothesis that cluster_2 arrays have spread between isolates because they provide immunity against MGEs that infect this population.

**Figure 5.**
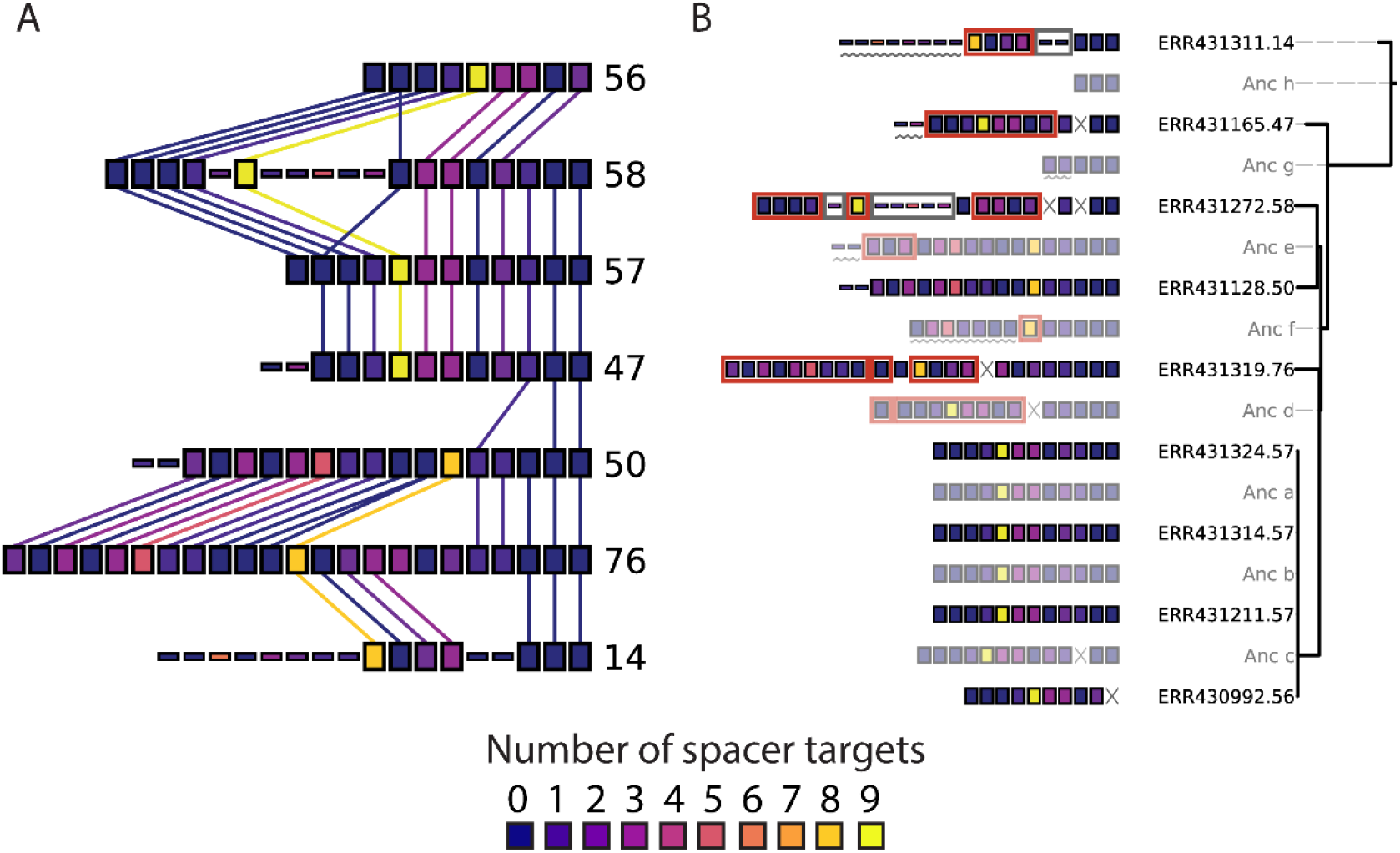
Spacerblast identifies protospacer targets of CRISPR spacers. Protospacer matches of spacers present in cluster_2 arrays were identified in assemblies using Spacerblast. Spacerblast was used with a requirement for 100% sequence identity between protospacer and spacer and the presence of a “CC” PAM 5’ of the protospacer sequence. Regions in which CRISPR arrays were identified were masked from the search using the Spacerblast “-r” option. Each spacer was assigned a colour according to the number of protospacers identified and these colour assignments were used as a custom colour scheme for CRISPRdiff (**A**) and Constrain (**B**).

### Assessing the performance of CRISPRtree using *in silico* evolved CRISPR arrays

As CRISPRtree uses a newly developed phylogenetic method, it is important to assess its ability to accurately infer the correct tree for a given set of arrays. However, we do not have access to a suitable dataset of CRISPR arrays with a known evolutionary history to test CRISPRtree. Therefore, we developed Evolve, a tool that generates simulated CRISPR arrays with known relationships through *in silico* evolution (Supplemental Methods). Evolve is distributed as part of CCTK to allow user assessment of CRISPRtree performance.

We used Evolve to generate a test dataset of CRISPR arrays that were generated with different evolution parameters (described in the Supplemental Methods). For each set of CRISPR arrays, Evolve stores the “true” tree describing the relationships between the simulated arrays. We then assessed the ability of CRISPRtree to infer the true tree when given only the arrays that were present at the end of each simulation (i.e., excluding any arrays that were “lost” during the *in silico* evolution process). Differences between the tree inferred by CRISPRtree and the true tree were measured using RF distance (See Methods). The performance of CRISPRtree for simulated datasets with different parameters is shown in **Figure 6**.

**Figure 6.**
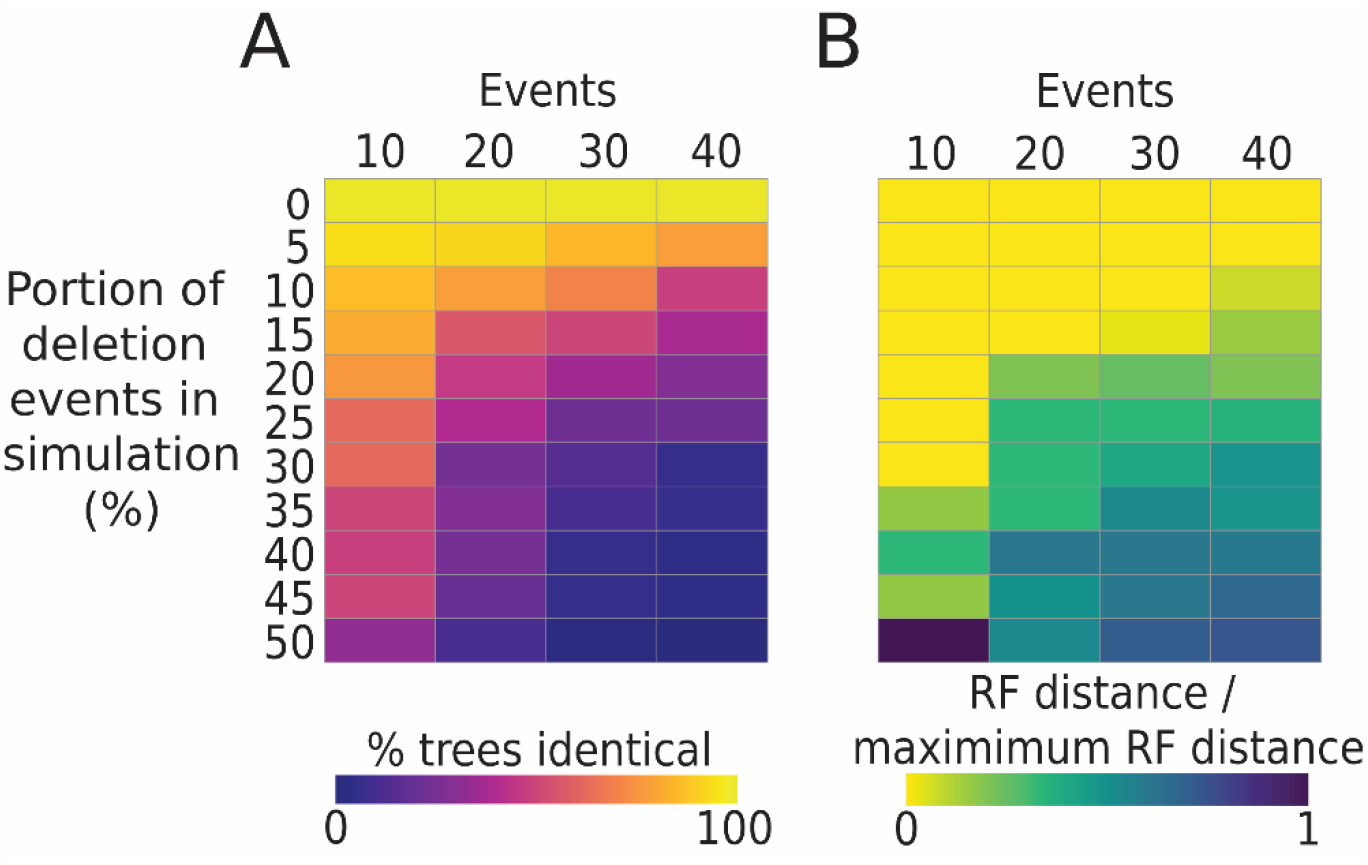
CRISPRtree can accurately recreate the true relationship between arrays when the frequency of spacer deletions is low. Simulated CRISPR arrays were produced using Evolve, and their relationships recorded in what will be referred to as the true tree topology. CRISPRtree was then used to analyze the simulated CRISPR arrays and to infer a maximum parsimony tree. The true topology of the simulated trees was compared to the topology inferred by CRISPRtree using Robinson-Foulds (RF) distance (Methods) ^29^. To allow the calculation of summary statistics between multiple sets of trees, RF distance is presented as a proportion of the maximum theoretical distance for each pair of trees. Simulations were performed with a range of parameters: different proportions of spacer acquisition and deletion events (rows within each heatmap); and number of events for which each simulation was run (columns within each heatmap). For each set of parameters, 50 replicate simulations were run and two summary statistics are reported: (**A**) the proportion of trees produced by CRISPRtree that are identical to the true tree topology (RF distance of 0); (**B**) the median ratio of the observed RF distance to maximum theoretical RF distance between the CRISPRtree and true tree topology.

When CRISPRtree was used to infer a tree for sets of arrays with known relationships, its performance was impacted by the frequency of deletions in the Evolve simulations (**Fig. 6**). If only spacer acquisitions occurred during the simulated evolution of CRISPR arrays, then CRISPRtree produced the correct tree every time regardless of the number of events (**Fig. 6A**; top row). However, when spacer deletions were included in the simulated CRISPR evolution, this resulted in a reduction in the accuracy of CRISPRtree in reconstructing the relationships between arrays (**Fig. 6A,B**; note darker colours in lower rows).

While small numbers of deletions reduced the ability of CRISPRtree to produce the correct tree, CRISPRtree was still able to produce a tree that is similar to the true tree for most of the parameter settings tested (**Fig. 6B**; light colours in most rows). Furthermore, for simulations involving lower numbers of events, and therefore producing arrays that are less diverged, CRISPRtree was able to produce the correct tree more than half the time even with high frequencies of deletions (**Fig. 6B**).

When the tree produced by CRISPRtree differed from the true tree, it was typically because the true relationships between arrays had been obscured by deletion events, resulting in arrays for which the most parsimonious tree is not the true tree. Examples of trees produced by CRISPRtree that differ from the true tree are presented in **Figure S4**. These trees show that CRISPRtree fails to identify the relationships between arrays in cases where deletions have removed evidence of those relationships.

Finally, we used the test dataset generated using Evolve to assess the impact of the parsimony cost of acquisitions, deletions, insertions, and independent acquisitions events on CRISPRtree performance (Supplemental Methods; **Fig. S5**). None of the tested event parsimony cost values improved CRISPRtree performance over the performance seen with default values. Instead, we found that CRISPRtree performance is robust to event cost. Specifically, CRISPRtree performance was equally good for all combinations of parsimony costs where the cost of acquisition is lowest, deletion is intermediate, and independent acquisition is highest. The parsimony cost of insertions had no impact on CRISPRtree performance in this analysis.

## Discussion

Here, we present CCTK - a toolkit for studying CRISPR arrays. CCTK includes tools to perform commonly used analyses, building on previously developed tools for the identification of CRISPR arrays (MinCED and BLASTN ^22,23^). In addition, we developed new resources for the analysis of CRISPR arrays (CRISPRdiff, CRISPRtree, Constrain and Spacerblast).

CRISPRdiff improves upon previously developed tools for the visualization of CRISPR arrays by providing a clearer visualization of array similarities and differences. CRISPRdiff also provides easier user control of the contents and appearance of the produced plot than existing tools and can be easily integrated into existing CRISPR-analysis pipelines.

CCTK also opens previously unexplored avenues of CRISPR research: CRISPRtree is the first tool developed for the inference of phylogenetic relationships between CRISPR arrays. CRISPRtree may be used to supplement other phylogenetic methods to further resolve relationships between highly related strains. Additionally, CRISPRtree enables researchers to reconstruct the immune history of coevolving virus-host partners both in laboratory experiments and in natural systems.

Constrain allows the reconciliation of CRISPR array relationships with other types of phylogenetic data and allows the detection of the HGT of whole CRISPR arrays or sets of spacers between isolates. Thus, Constrain can improve our understanding of the role of CRISPR array HGT in the microbial pan-immune system ^40^.

By combining the above tools into a single command-line interface, CCTK makes the analysis of CRISPR arrays easy and scalable and is a valuable new resource for the study of CRISPR systems.

## Acknowledgements

The authors would like to thank Tandy Warnow and Bernard Moret for helpful discussions about the phylogenetic approaches used in this study. The authors are also grateful to Giulia Orazi for critical reading of the manuscript.

## Figure legends

**Figure S1. Phylogenetic relationships between isolates analyzed in this study**

Sequence records were retrieved for the list of “clone corrected” isolates described by England et al. ^16^. Assemblies were generated using Spades with the “careful” option ^18^. Each leaf label corresponds to the European Nucleotide Archive accession number of the reads used to generate that assembly. Core genome SNPs were identified using Spine and Nucmer and aligned using a custom script (See Supplemental Methods). IQTREE2 was then used to infer a maximum likelihood tree using the core genome alignment and a model determined using the ModelFinder utiltity (model selected: TVM+F+R4) ^21,43^. Bootstrap support was calculated using the Ultrafast Bootstrap utility which recommends a threshold of 95% support for considering a node to be supported (Red nodes not supported) ^37^. The tree was visualized using iToL, midpoint rooted, and annotated according to whether any CRISPR arrays were identified (inner, black ring) and the presence of arrays belonging to the largest 7 clusters shown in Figure 2 (coloured rings; see key) ^44^. Isolates encoding arrays from cluster_2 are indicated both with a blue ring and with a blue leaf label background.

**Figure S2. The process used by CRISPRtree to infer a tree explaining CRISPR array relationships**

CRISPRtree infers a tree using the following process: (**A**) CRISPRtree starts with a list of arrays to analyze. (**B**) The first and second array in the list are aligned and modules of spacer in their alignment are identified (indicated using coloured bars beneath the aligned arrays). A module of spacers is a group of consecutive spacers that have the same kind of relationship between the two arrays. For example, the indel module indicated by a green bar in the middle of the aligned arrays corresponds to a set of consecutive spacers that are present in one array but not in the other. (**C**) A hypothetical ancestral array is formed. Each module is then processed to decide which spacers would be present in the ancestral array (See Supplemental Methods). (**D**) A tree is initialized with the two extant arrays and their ancestor. (**E**) The next array in the list is aligned to each array already present in the tree (Arrays 1, 2, and Anc_a). Modules are identified based on each alignment and the parsimony cost of each event indicated by the modules is calculated. The cost is calculated for both arrays being aligned and the greater of the two costs is taken as the score for that alignment. e.g., for the alignment of array 3 and 2, array 3 is found to have 3 acquired spacers and an indel with default parsimony cost of 3+30=33. Array 2 is found to have 8 acquired spacers, an indel, and a trailer loss with default parsimony cost of 8+30+1=39. Therefore, the parsimony cost of this alignment is 39. Array 3 when aligned with either array 1 or Anc_a has the same parsimony cost. In that case the ancestral array is preferred. (**F**) A hypothetical ancestral array is inferred for arrays 3 and Anc_a using the same process as shown in panels **B** and **C**. (**G**) Array 3 and its ancestor, Anc_b, are added to the tree. (**H**) Once all arrays in the initial list have been added to the tree, events are annotated on each array. The parsimony cost of all annotated events is used as a score for the tree. This score can be compared between trees to identify the most parsimonious tree. Different trees are produced by using a different order of arrays in the initial list.

**Figure S3. CRISPRdiff produces a colourblind-friendly visualization while CRISPRStudio and CRISPRviz visualizations have low visual contrast**.

The arrays shown in Figure 3 are shown with the two most common forms of colourblindness simulated using Adobe Illustrator.

**Figure S4. CRISPRtree fails to infer the correct tree when arrays are degraded through deletion**.

Three examples in which CRISPRtree did not accurately identify the relationships between simulated arrays. In each panel, the tree on the left represents the true relationships of simulated arrays produced by Evolve, while the tree on the right was inferred by CRISPRtree. For each example (separate panel), arrays that have been misplaced by CRISPRtree are indicated with asterisks adjacent to the array number. Each example corresponds to a different set of simulation parameters: (**A**) 20 events, 65% acquisition, 35% deletion, 85% loss rate, (**B**) 40 events, 60% acquisition, 40% deletion, 75% loss rate, (**C**), 40 events, 70% acquisition, 30% deletion, 85% loss rate. Beneath each pair of trees, the ratio is shown of the Robinson-Foulds (RF) distance / the maximum RF distance. In the true trees, ancestral arrays are numbered according to which event number they were created by during the Evolve simulation.

**Figure S5. Impact of event parsimony cost on accuracy of CRISPRtree in reconstructing true tree topology**.

(**A-D**) Relative performance of CRISPRtree assessed for simulated data with different loss rates: (**A**) 75%, (**B**) 80%, (**C**) 85%, (**D**) 90%. (**E**) Relative performance of CRISPRtree using an acquisition parsimony cost of 10 when analyzing data generated with a loss rate of 75% (compare to panel **A** in which loss rate is the 75% and acquisition cost is 1). In each panel (**A-D**), the heatmap corresponding to default parameters is omitted. (**F**) The mean value of each heatmap shown in panels **A-E** is summarized as a single value. Shown is the impact of the parsimony cost of deletions (columns of each heatmap) and independent acquisitions (rows of each heatmap) on CRISPRtree performance. Only heatmaps for insertion cost of 30 were summarized here.

