## Supplemental Methods and Figures for "CRISPR comparison toolkit (CCTK): Rapid identification, visualization, and analysis of CRISPR array diversity"

#### CRISPR identification by CCTK Minced

To identify CRISPR arrays using CCTK Minced [1], CCTK first runs MinCED using default settings on each input assembly. The output files generated by MinCED are then processed to extract the location of each predicted CRISPR array as well as repeat and spacer sequences.

Each array in the MinCED output files is processed as follows. First, the number of occurrences of each repeat sequence variant in the array (if variations exist in the repeat sequence) is counted and the most common is retained and considered the representative repeat associated with that array. This repeat is then compared to either a user-provided database of CRISPR repeat sequences or to a built-in database of repeat sequences (types I-A, I-C, I-E, and I-F) for which the correct orientation relative to the array leader end is known. Finally, the best-matching repeat in the database is used to orient the spacers in the array and the CRISPR subtype is added to the identifiers of all spacers in this array.

The process by which a repeat is compared to the database of repeats is as follows. The Hamming distance between the query repeat and each database repeat is calculated for both the sequence found by MinCED and the reverse complement of that sequence. The database repeat with the lowest Hamming distance is considered the best match and, if the lowest Hamming distance was found when the reverse complement of the query repeat was considered, then the array is reversed (i.e., each spacer is converted to the reverse complement sequence and the order of spacers is reversed.) The CRISPR type associated with the best matching repeat in the database (the fasta header if using a user-provided database) is added to the IDs for each spacer in that array. However, if the Hamming distance between query repeat and database repeat is greater than 5, the best match repeat is still used to orient the array, but each spacer is assigned an ID of "unknown_CRISPR_type" and the CRISPR type of the best matching database repeat is included in parentheses. Poorly matching repeats are still used to orient arrays and "unknown_CRISPR_type" IDs assigned so that CRISPR repeats that do not match any repeats are conspicuous, but if there are multiple arrays of the same unknown type, they will be consistently oriented.

#### CRISPR identification by CCTK Blast

To identify CRISPR arrays using CCTK Blast [2], CCTK requires the user to provide a fasta format file containing all repeats for the CRISPR types to search for, and a blast database made using makeblastdb with the "-parse_seqids" option. If the user is searching for CRISPR arrays in a blast database that contains assemblies that are each in multiple contigs, then the user can specify a file containing an identifier that is present in all contigs associated with the same assembly. Alternatively, the user can describe this identifier using a regular expression. If the assemblies are in multiple contigs but such an identifier is not described then the arrays identified by CCTK will be unaffected, but each contig will be treated as a separate assembly when reporting which arrays were found in which assemblies.

Arrays are identified in the provided BLAST database by searching for the provided repeat sequences and processing the BLASTN output. The command used to search for repeats (using default cctk blast settings) is "blastn -query <user repeats> -db <user database> -task blastn-short -outfmt '6 std qlen slen sseq' -num_threads 1 -max_target_seqs 10000 -evalue 10". Each result is then checked to see if the

BLAST hit covered the entire length of the query sequence. If it did not, then the hit is extended to cover the full length of the query by retrieving the missing sequence using blastdbcmd. The percent identity between each hit and the corresponding query is then calculated and those that do not meet the user-specified cutoff (default 80%) are removed from further consideration.

Remaining BLAST hits (i.e., full length matches with over 80% identity) are then sorted according to the contig and position at which they were identified. BLAST hits are grouped into arrays by working through the ordered list of BLAST results and grouping consecutive hits together if they meet the following criteria: fewer than a user-specified number of bases apart (measured from start of a hit to start of the next hit; default = 80), on the same strand of DNA, in the same contig. Note that it is not possible to search for repeats with highly similar sequence using this method as the blast hits for each variant repeat will overlap. Groups of more than three repeats that meet these criteria are considered to be part of the same CRISPR array. Spacer sequences between each repeat in the array are then extracted using blastdbcmd. The strand from which spacer sequence is extracted is determined by the direction in which the user-provided repeat sequence matched. Spacers are assigned IDs containing the fasta header of the repeat in the user-provided file with which they are associated.

### Deduplicating similar CRISPR spacers

Once arrays have been identified, spacers that differ by fewer than a user-specified threshold (similar spacers) can be deduplicated, leaving a single representative of each set of similar spacers. This is achieved through the following process. First, similar spacers are identified by using BLASTN with the entire set of spacers as both the query and subject sequences. BLASTN is used because performing all pairwise comparisons is infeasible when working with large numbers of spacers. The results of this BLAST are then processed. If the number of mismatches reported by BLAST exceed the user-specified cutoff, then the result are discarded. If the length of the match returned by BLAST is not the full length of both spacers, then the sequence of the two spacers is compared using Hamming distance. This is done for both unmodified sequences, and with one or the other sequence shifted one position to catch examples of otherwise identical spacers that differ in length by a single base. If the number of mismatches reported by BLAST for full length hits, or the Hamming distance of other hits is equal to or less than the user-specified cutoff, then the hits are processed further.

Next, similar spacers are represented as a network to identify clusters of related spacers. In this network spacers are represented as nodes and an edge is drawn between them if they differ by fewer than the mismatch cutoff. A set of spacers that are connected to one another by any number of intermediate edges is then considered a cluster (i.e., even if two spacers are only connected via an intermediate spacer, they are considered part of the same cluster). Representatives are then chosen for each cluster as follows. First, any clusters that are completely connected (i.e., all spacers in the cluster are connected by an edge to all other spacers) are reduced to a single representative. The representative is chosen by counting the number of occurrences of each cluster member in the dataset of identified spacers. The most numerous spacer is chosen as the representative. Clusters that are not completely connected are then processed by first breaking them into smaller clusters. This is done to avoid considering spacers that differ by more mismatches than the user-specified threshold to be the same spacer. First, the number of edges connected to each spacer in the cluster is counted. The most connected spacer and all spacers with which it shares an edge are then removed from the original cluster and considered to be a new cluster. This process is repeated with the remaining spacers in the original cluster until no spacers remain in the original cluster. Then, each of the spacers identified as the most connected in each newly formed cluster is chosen as the representative of that cluster. After each set of similar spacers has been processed and a representative chosen, the sequence and ID of the representative is used to replace all other cluster members in the dataset. Information about which sequences were identified as similar to each representative are then written to a file in the output directory.

### CRISPRdiff: CRISPR array colour assignment, order determination, and plotting

Spacer colour assignments are performed through two steps. First, the set of spacers that are found in more than one array is identified and assigned a colour. Spacer colour-assignment is performed as follows:

- When fewer than 64 different spacers must be assigned a colour, a colour blind-friendly colour scheme is used [3].
- When over 64 spacers are to be assigned a colour, the first 64 are assigned colours from the preset colour scheme and the remainder are assigned random colours.
- Any spacers found in only a single array is represented as a thin, black rectangle.
- Colours can also be provided by the user and can be stored in JSON format and used by other CCTK tools to allow consistent colour assignments to spacers between images produced by CCTK tools.

Array order can be determined by three methods depending on user-settings:

- The user may set the order to be used.
- Alternatively, the user may set what they think is approximately the correct order (for example if two obvious clusters of arrays are being visualized). CRISPRdiff then uses a simple search to improve the order according to the following process:
    - The number of spacers shared between neighbouring arrays is counted and considered the score of that order.
    - Then, the position of the first and second arrays is swapped, and the score is recalculated. If the score has increased, then the new order is kept. If not, then the order is reset.
    - Next, the position of the second and third arrays is swapped and so on until the end of the list. If any changes result in an improvement in the score, then the process is repeated from the beginning to the end of the list again.
    - This process is repeated through the entire list of arrays until the list is traversed without improving the score.
- If the user does not specify an array order, then for small numbers of arrays (fewer than nine) all possible array orders are checked and the best is used.
- For larger numbers of arrays, the following process is used to find a good order:
    - First, the order of arrays in the list is shuffled and the order is scored according to the number of spacers shared between all pairs of neighbouring arrays.
    - Next, the position of a random array is swapped with that of another randomly selected array and the score recalculated. If the score increased, the new order is kept. If not, the order is reset.

- Then, another pair of randomly selected arrays are swapped and the process repeated. This process is repeated until 100 swaps do not result in an improvement in the score.
- In addition, this entire process is repeated starting with a different random initial order of arrays for a number of replicates that can be set by the user. The order that produces the highest score among all replicates is used for plotting.

Once the array order has been chosen, arrays are visualized using Matplotlib [4]. Spacers are drawn as squares for which the fill and outline colour combination is unique to each spacer. Lines are then drawn connecting identical spacers present in adjacently plotted arrays. These lines are assigned the same colour as the spacer being connected.

### CRISPRtree inference of ancestral arrays

Starting from the trailer end of the arrays aligned in **Figure S2B**, the indicated modules would be processed as follows:

- Trailer-loss modules: the lost spacer is added to the ancestor.
- Shared modules: all shared spacers are added the ancestor.
- Indel modules: Indels are processed differently depending on the specific relationship.
    - If one array has spacers in the module, while the other has none, then a deletion is inferred to have occurred and all the spacers are added to the ancestral array
    - If both arrays have different spacers in this module (i.e., a mismatch rather than gap in the alignment), then the other arrays being analyzed are used to resolve this event (i.e., not the two being compared here). First, all arrays are searched for the two sets of spacers in the module.
        - If neither set of spacers in the module being assessed is found, then one set is chosen at random to add to the ancestor (No evidence found to support the choice of one set of spacers over the other, but one insertion is more parsimonious than two).
        - If only one set is found, then that set is added to the ancestor and the other is not.
        - If both sets are found, then both sets are added to the ancestor. The order of the two sets of arrays is determined by searching the arrays in the dataset. If the two sets of arrays are not seen together as a consecutive run of spacers in any array, then a random order of the two sets is chosen.
- Acquisition modules: differences at the leader end are assumed to have happened since the ancestor so these spacers are not added to the ancestor.

### CRISPRtree and Constrain identification of independent acquisitions of spacers

Once all arrays have been added to a tree, all leader-end acquisition and insertion events are reconsidered to identify independent acquisitions. This is achieved by considering each acquisition or insertion module one at a time (in both ancestral and extant arrays). When considering the modules in an array, the process is as follows:

- First, all arrays in the tree are divided into two groups: arrays that are descendants of the array being considered (referred to here as “are-descendants group”), and those that are not (“not-descendants group”). As the modules being considered here are insertion and acquisition modules, and therefore contain spacers that were not present in their direct ancestor, the presence of spacers in arrays in the not-descendants group may indicate independent acquisition.

- Next, the spacers in the module are compared to the spacers in all arrays that are in the not-descendants group. If no matching spacers are found, then no independent acquisition is inferred. However, if matching spacers are found, then the following steps are followed to classify spacers into independent acquisition events.
    - If all spacers in the module are identified as a single consecutive set of spacers in one or more arrays in the not-descendants group, then the whole module is classified as an independent acquisition.
    - If only some spacers are found as a single consecutive set of spacers in one or more arrays in the not-descendants group, then the matching spacers are classified as an independent acquisition event. Spacers from the module that were not found retain their original classification. If the newly classified independent acquisition module is in the middle of an existing module, this results in the existing module becoming two modules of the original type either side of a newly formed independent acquisition module.
    - If different combinations of the spacers are found in multiple different arrays in the not-descendent group, then the module is processed using a repeating process as follows. First, the longest stretch of consecutive spacers found in an array in the not-descendant group is identified. This stretch of spacers is classified as an independent acquisition module. The process then repeats with the remaining spacers that have not been classified until all spacers have been classified or until no more spacers are found in the not-descendant group.

Whenever a set of spacers is identified as independently acquired, all the arrays in the not-descendants group that contain the set of spacers is reported to the user in the log. This information is also annotated on the tree image produced by CRISPRtree or Constrain.

### Spacerblast: identification of protospacers and protospacer adjacent motifs

Spacerblast identifies possible protospacers by searching for CRISPR spacer matches in a BLAST database using BLASTN and blastdbcmd [2]. This is achieved through three steps.

- First, initial matches are identified using BLASTN using the following command (when Spacerblast is run with default settings) “blastn -query <spacers> -db <user database> -task blastn-short -outfmt '6 std qlen slen' -num_threads 1 -max_target_seqs 10000 -evalue 10”.
- Next, any BLAST results that do not cover the full length of the query spacer are extended by retrieving the missing sequence using blastdbcmd. The percent identity between query spacers and full length matches is then assessed and those which do not meet the user-specified cutoff (default: no cutoff) are discarded.
- Finally, if the user has specified a protospacer-adjacent motif (PAM) then sequence flanking the match is retrieved using blastdbcmd. The user can specify the location of the PAM either 5’ or 3’ of the protospacer and can define the PAM sequence using either IUPAC nucleotide codes [5] or a regular expression.

The sequence flanking each BLAST match is then checked for the presence of the described PAM and BLAST matches are sorted based on whether there is a PAM or not. Results are then returned to the user. By default, only BLAST matches with a flanking PAM are returned. However, matches without a flanking PAM can also be saved to a file specified by the user.

### Evolve: *in silico* evolution of CRISPR arrays

Each simulation starts with a single array of 5 spacers and proceeds for a fixed number of events, during which an array is selected at random and duplicated. The copy of the original array is then modified either by the acquisition of a new spacer at its leader end, a deletion of one or more spacers from any location, or loss of a single spacer from the trailer end. The choice of acquisition, deletion, or trailer-loss is made according to probabilities that are set for each simulation. After the duplicated array is modified, the original array may be removed from future consideration based on a probability referred to as “loss rate.” If the array is removed, it can no longer be selected for future events.

Deletion and trailer-loss events are only applied to arrays with a length greater than 1 (i.e., if an array of length 1 is being processed and a deletion is chosen from the possible events, an acquisition is chosen instead). Deletions are applied using the following process.

- First two positions in the array are chosen by sampling from a normal distribution generated using NumPy. This normal distribution has a mean equal to the length of the array divided by 2 and a standard deviation equal to the length of the array divided by 4 [6].
- If the two numbers selected are identical or if the numbers represent the start and end indices of the array, then one of the numbers is chosen again until those conditions are no longer met.
- Next, all spacers in the range of indices between the two numbers are removed from the array.

The use of a normal distribution centered on the middle of the array is intended to approximate the greater probability of a spacer being lost in the middle of the array. This bias is due to a larger number of repeat sequences either side of centrally-located spacers that could take part in a recombination event with a different repeat. The relationship between spacer position in an array and probability of deletion has been proposed previously [7].

### Core genome SNP identification using Spine and Nucmer

To generate an alignment of core genome SNPs, Spine is used to identify the core segments of the set of assemblies being considered [8]. Next, Nucmer is used to align the core genome segments identified by Spine [9]. Nucmer is also used to identify SNPs between unambiguously aligned core segments (i.e., segments that only align in a single location). Finally, a custom python script (snps2fasta.py; available at GitHub URL below) is used to extract SNP locations and sequence from Nucmer output files and the core genome fasta file produced by Spine. The scripts used to process Spine and Nucmer outputs into a core genome SNP alignment are provided here https://github.com/Alan-Collins/Spine-Nucmer-SNPs. In addition, a full description of how to run a pipeline of Spine, Nucmer, and the scripts used here to make a core genome SNP alignment is described in the readme of that GitHub repo. These steps are described in more detail below.

To produce the trees used in this study, the following specific process was used. First Spine (version 0.3.2) was run with default parameters. Spine produces a core genome multifasta for each input assembly as well as a file (output.backbone.fasta) which will be referred to here as the “reference core”.

Next, Nucmer (version 3.1) was used to align the core genome contigs identified by Spine for each assembly to the reference core, followed by the delta-filter utility included with Nucmer using the -r and -q options and then show-snps utility with the options -C -l and -r. Finally, snps2fasta.py was run using -whole to produce an alignment of the entire core genome. The process by which snps2fasta.py works is described below.

The script snps2fasta.py produces a core genome alignment using the following process. First, each of the .snps files produced by the MUMmer utility show-snps are read. Those files contain SNP information including the location in the reference core, the base in the reference core, and the base in the sequence being aligned to the reference core. Only positions that differ between the sequence being aligned and the reference core are present in each file. Therefore, as the files are read, a list of the positions that varied in any of the aligned sequences is constructed. For each sequence, the SNP base is stored if the sequence differed at a position. If the sequence did not differ at a position, then the base at that position in the reference core is used.

In addition to SNPs, Nucmer also identifies indels (insertion/deletion events). When these are encountered, all the aligned sequences lacking bases at the indel position are adjusted with dash symbols.

If the user has specified that they want the entire core genome alignment using the -whole option then an additional step is performed. Once all variant positions have been identified and either variant or reference core sequence has been added to each sequence being aligned, the remaining invariant sequence for the entire reference core is filled in. The aligned sequences is then output as a single concatenated sequence for each core assembly that was aligned to the reference assembly. Optionally, the sequences can be output as a separate alignment file for each contig present in the reference core, and the SNP information can be output in a table format in which row names are the name of each core assembly being aligned and column names are the location of each SNP relative to the reference core.

### Assessment of the impact of event parsimony cost on CRISPRtree performance

CRISPR arrays generated by Evolve were analyzed using CRISPRtree with different parsimony costs for acquisitions, deletions, insertions, and independent acquisitions. For each set of parameters, the performance of CRISPRtree was assessed relative to the performance of CRISPRtree when run with the default parsimony costs (as presented in **Figure 6**). Performance was assessed both in terms of the number of trees produced by CRISPRtree that were identical to the “true” tree (percent of trees identical) and the extent to which trees produced by CRISPRtree differ from the true tree (Robinson-Foulds (RF) value between CRISPRtree and true tree divided by the maximum theoretical RF distance between those tree topologies; median RF ratio). Each heatmap in Figure **S5** represents the performance of CRISPRtree for arrays generated with the same evolution parameters as those shown in **Figure 6** (i.e., the 4 columns of each heatmap correspond to the number of events in the simulation, while rows correspond to the frequency of deletion events). The performance of CRISPRtree with the indicated parsimony costs is shown relative to the default parsimony costs. Performance is scaled such that for the “percent of trees identical” measure, the worst performance indicates that the tested values produced no identical trees while default values produced only identical trees. For the “median RF ratio” measure, the worst performance indicates that the tested values produced trees that were as different as theoretically possible from the true tree, while CRISPRtree produced only identical trees when run with default values.

### Supplemental figures

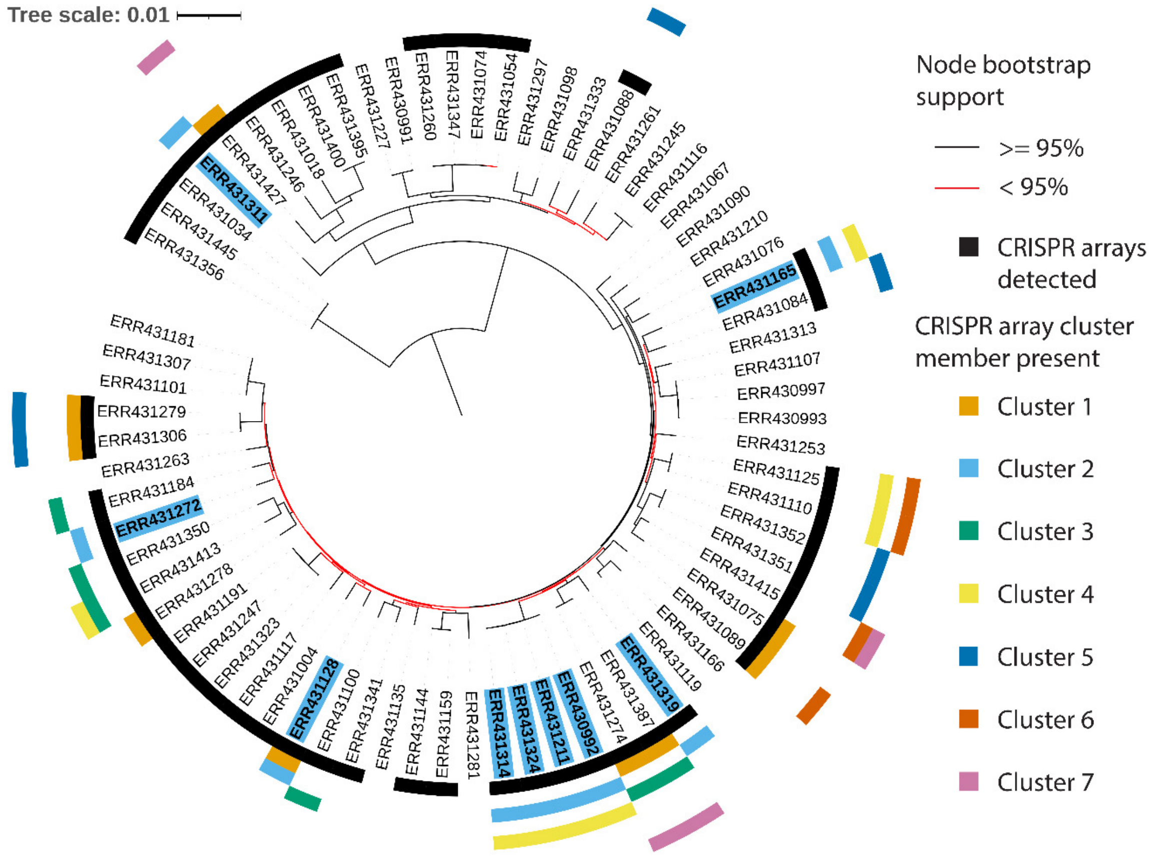


**Figure S1. Phylogenetic relationships between isolates analyzed in this study.** Sequence records were retrieved for the list of “clone corrected” isolates described by England et al. [10]. Assemblies were generated using Spades with the “careful” option [11]. Each leaf label corresponds to the European Nucleotide Archive accession number of the reads used to generate that assembly. Core genome SNPs were identified using Spine and Nucmer and aligned using a custom script (See Supplemental Methods). IQTREE2 was then used to infer a maximum likelihood tree using the core genome alignment and a model determined using the ModelFinder utiltity (model selected: TVM+F+R4) [12,13]. Bootstrap support was calculated using the Ultrafast Bootstrap utility which recommends a threshold of 95% support for considering a node to be supported (Red nodes not supported) [14]. The tree was visualized using iToL, midpoint rooted, and annotated according to whether any CRISPR arrays were identified (inner, black ring) and the presence of arrays belonging to the largest 7 clusters shown in Figure 2 (coloured rings; see key) [15]. Isolates encoding arrays from cluster_2 are indicated both with a blue ring and with a blue leaf label background.

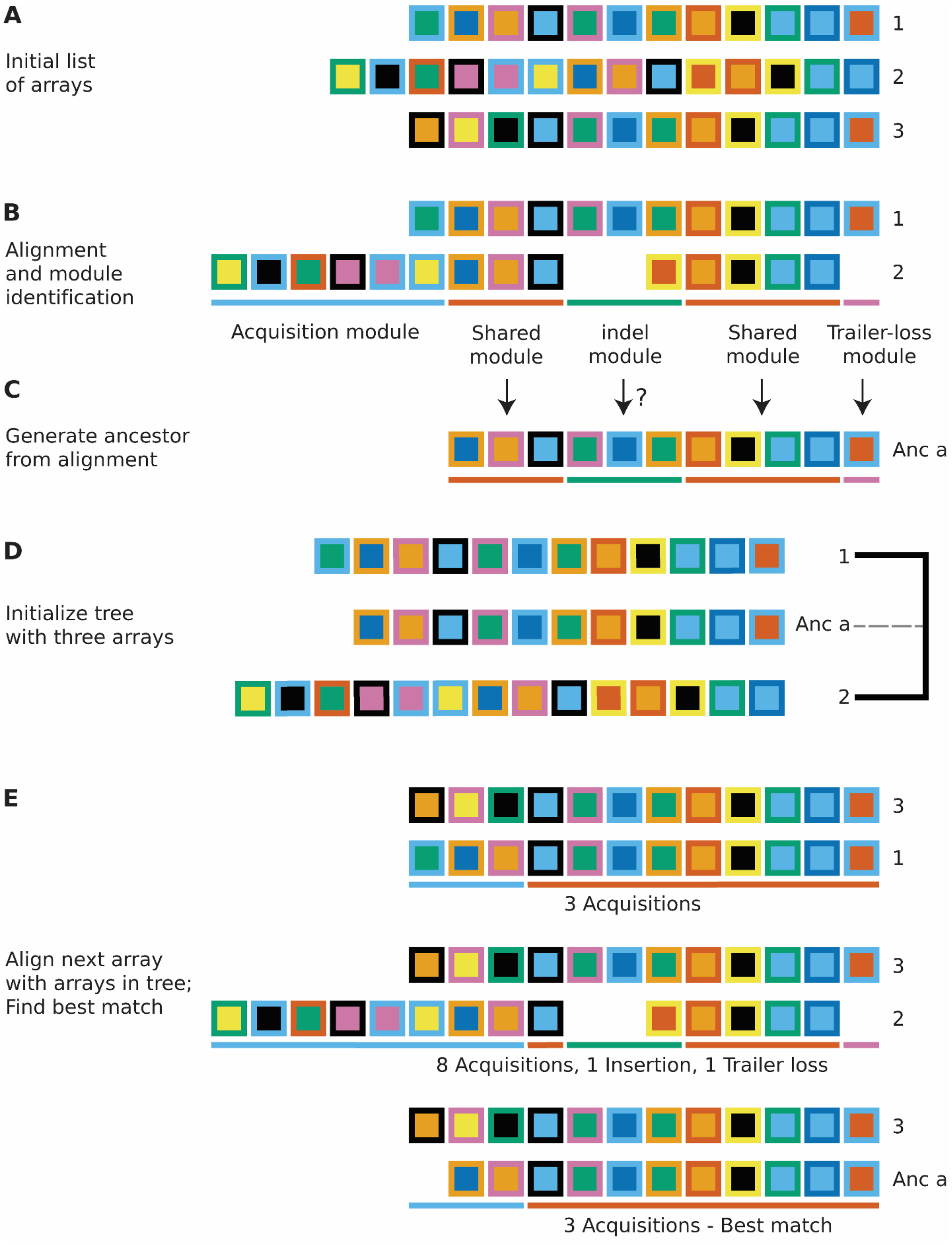
A
Initial list of arrays
1
2
3
B
Alignment and module identification
1
2
Acquisition module
Shared module
indel module
Shared module
Trailer-loss module
C
?
Generate ancestor from alignment
Anc a
D
Initialize tree with three arrays
1
Anc a
2
E
3
1
3 Acquisitions
Align next array with arrays in tree; Find best match
3
2
8 Acquisitions, 1 Insertion, 1 Trailer loss
3
Anc a
3 Acquisitions - Best match

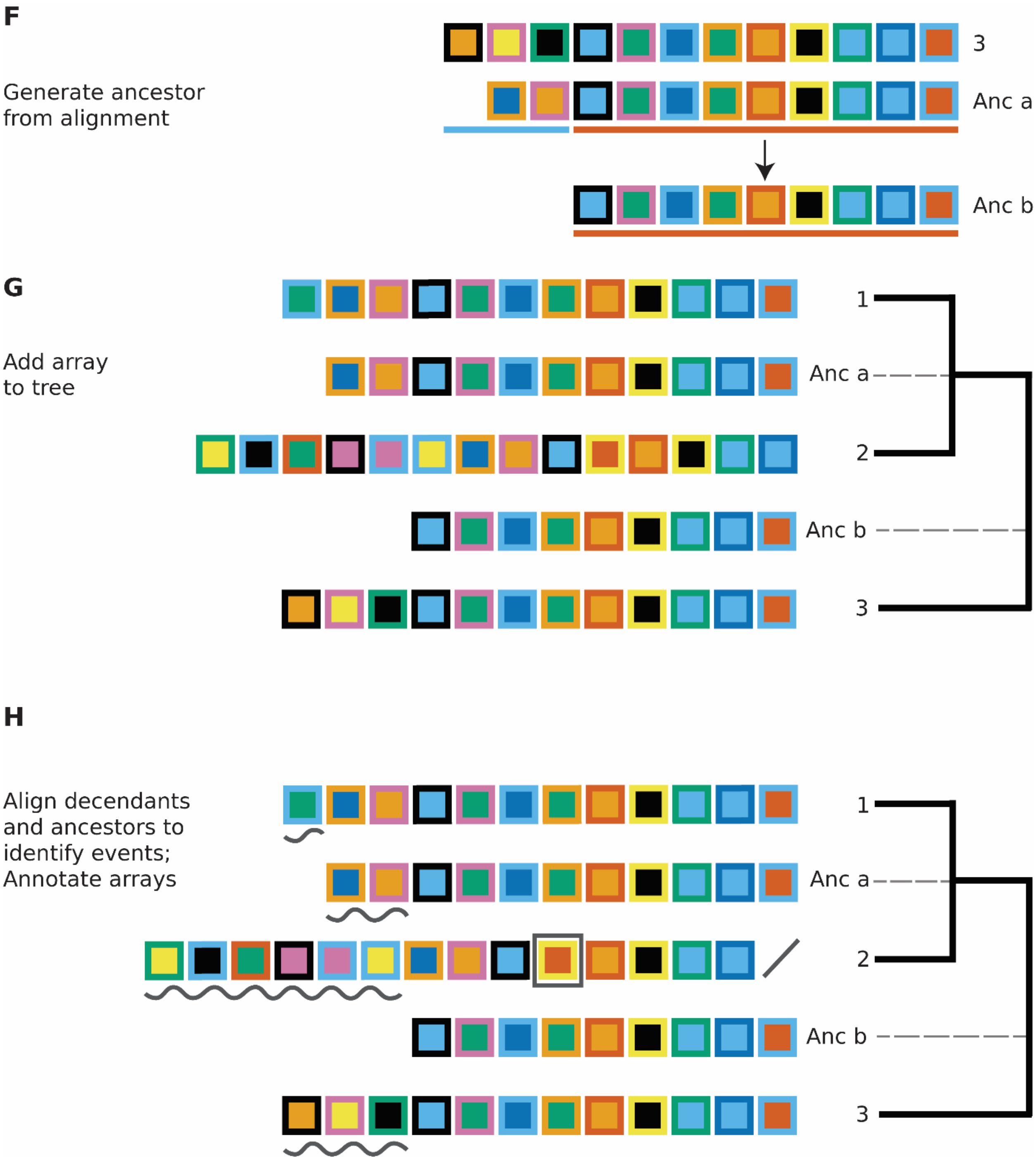


**Figure S2. The process used by CRISPRtree to infer a tree explaining CRISPR array relationships.** CRISPRtree infers a tree using the following process: (**A**) CRISPRtree starts with a list of arrays to analyze. (**B**) The first and second array in the list are aligned and modules of spacer in their alignment are identified (indicated using coloured bars beneath the aligned arrays). A module of spacers is a group of consecutive spacers that have the same kind of relationship between the two arrays. For example, the indel module indicated by a green bar in the middle of the aligned arrays corresponds to a set of consecutive spacers that are present in one array but not in the other. (**C**) A hypothetical ancestral array is formed. Each module is then processed to decide which spacers would be present in the ancestral array (See Supplemental Methods). (**D**) A tree is initialized with the two extant arrays and their ancestor. (**E**) The next array in the list is aligned to each array already present in the tree (Arrays 1, 2, and Anc_a). Modules are identified based on each alignment and the parsimony cost of each event indicated by the modules is calculated. The cost is calculated for both arrays being aligned and the greater of the two costs is taken as the score for that alignment. e.g., for the alignment of array 3 and 2, array 3 is found to have 3 acquired spacers and an indel with default parsimony cost of 3+30=33. Array 2 is found to have 8 acquired spacers, an indel, and a trailer loss with default parsimony cost of 8+30+1=39. Therefore, the parsimony cost of this alignment is 39. Array 3 when aligned with either array 1 or Anc_a has the same parsimony cost. In that case the ancestral array is preferred. (**F**) A hypothetical ancestral array is inferred for arrays 3 and Anc_a using the same process as shown in panels **B** and **C**. (**G**) Array 3 and its ancestor, Anc_b, are added to the tree. (**H**) Once all arrays in the initial list have been added to the tree, events are annotated on each array. The parsimony cost of all annotated events is used as a score for the tree. This score can be compared between trees to identify the most parsimonious tree. Different trees are produced by using a different order of arrays in the initial list.

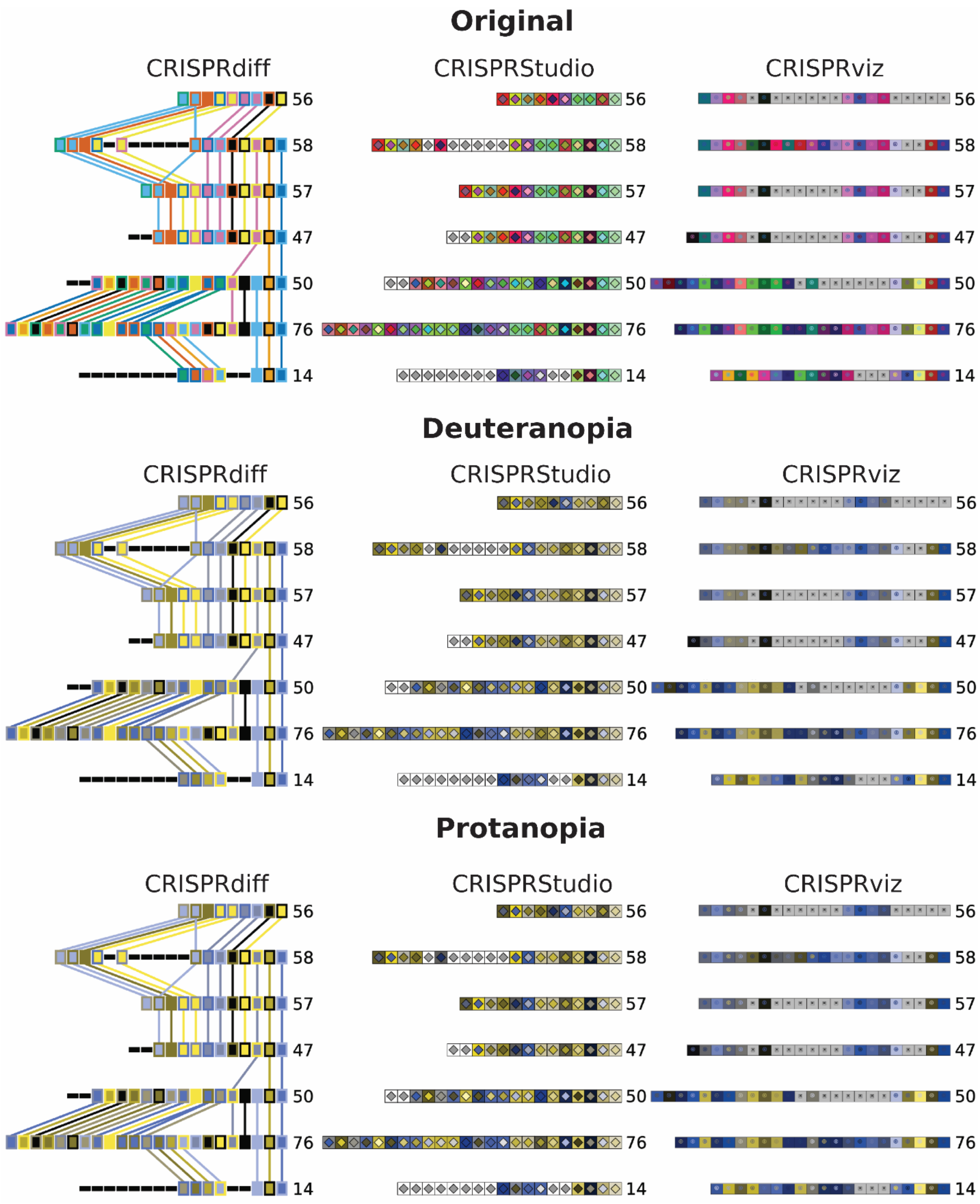


**Figure S3. CRISPRdiff produces a colourblind-friendly visualization while CRISPRStudio and CRISPRviz visualizations have low visual contrast.** The arrays shown in Figure 3 are shown with the two most common forms of colourblindness simulated using Adobe Illustrator.

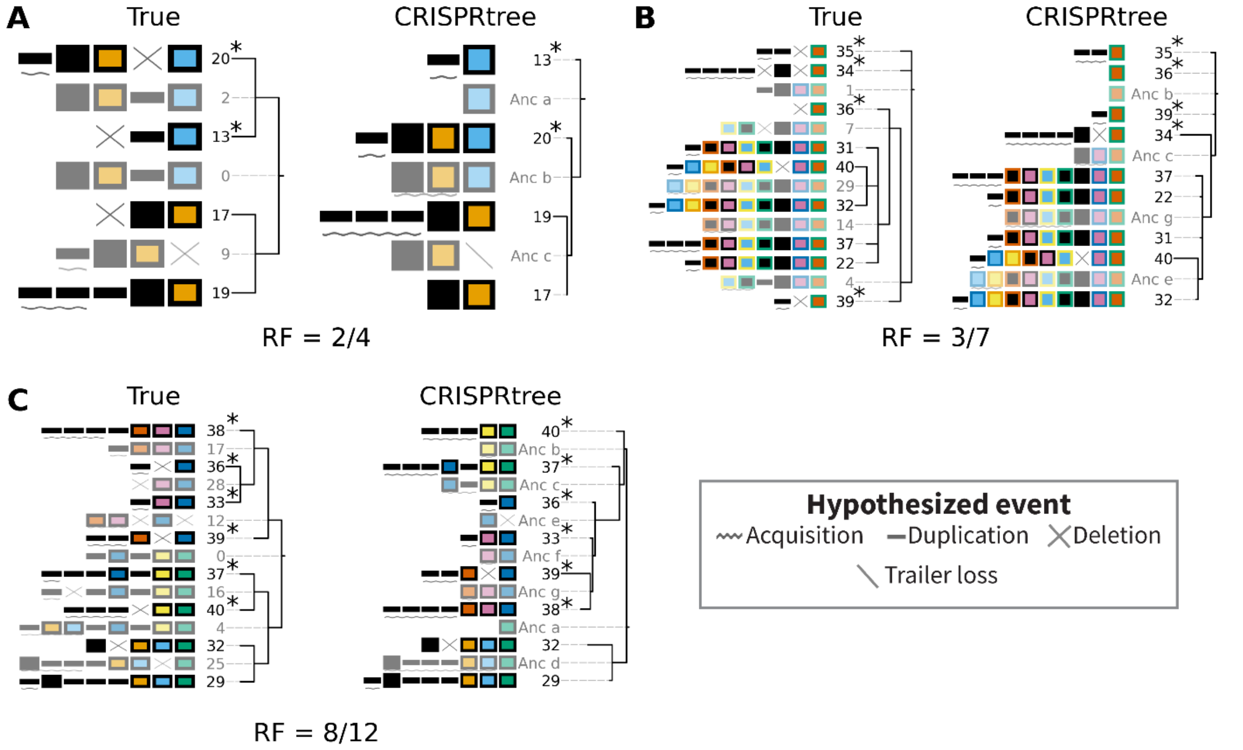


**Figure S4. CRISPRtree fails to infer the correct tree when arrays are degraded through deletion.** Three examples in which CRISPRtree did not accurately identify the relationships between simulated arrays. In each panel, the tree on the left represents the true relationships of simulated arrays produced by Evolve, while the tree on the right was inferred by CRISPRtree. For each example (separate panel), arrays that have been misplaced by CRISPRtree are indicated with asterisks adjacent to the array number. Each example corresponds to a different set of simulation parameters: (**A**) 20 events, 65% acquisition, 35% deletion, 85% loss rate, (**B**) 40 events, 60% acquisition, 40% deletion, 75% loss rate, (**C**), 40 events, 70% acquisition, 30% deletion, 85% loss rate. Beneath each pair of trees, the ratio is shown of the Robinson-Foulds (RF) distance / the maximum RF distance. In the true trees, ancestral arrays are numbered according to which event number they were created by during the Evolve simulation.

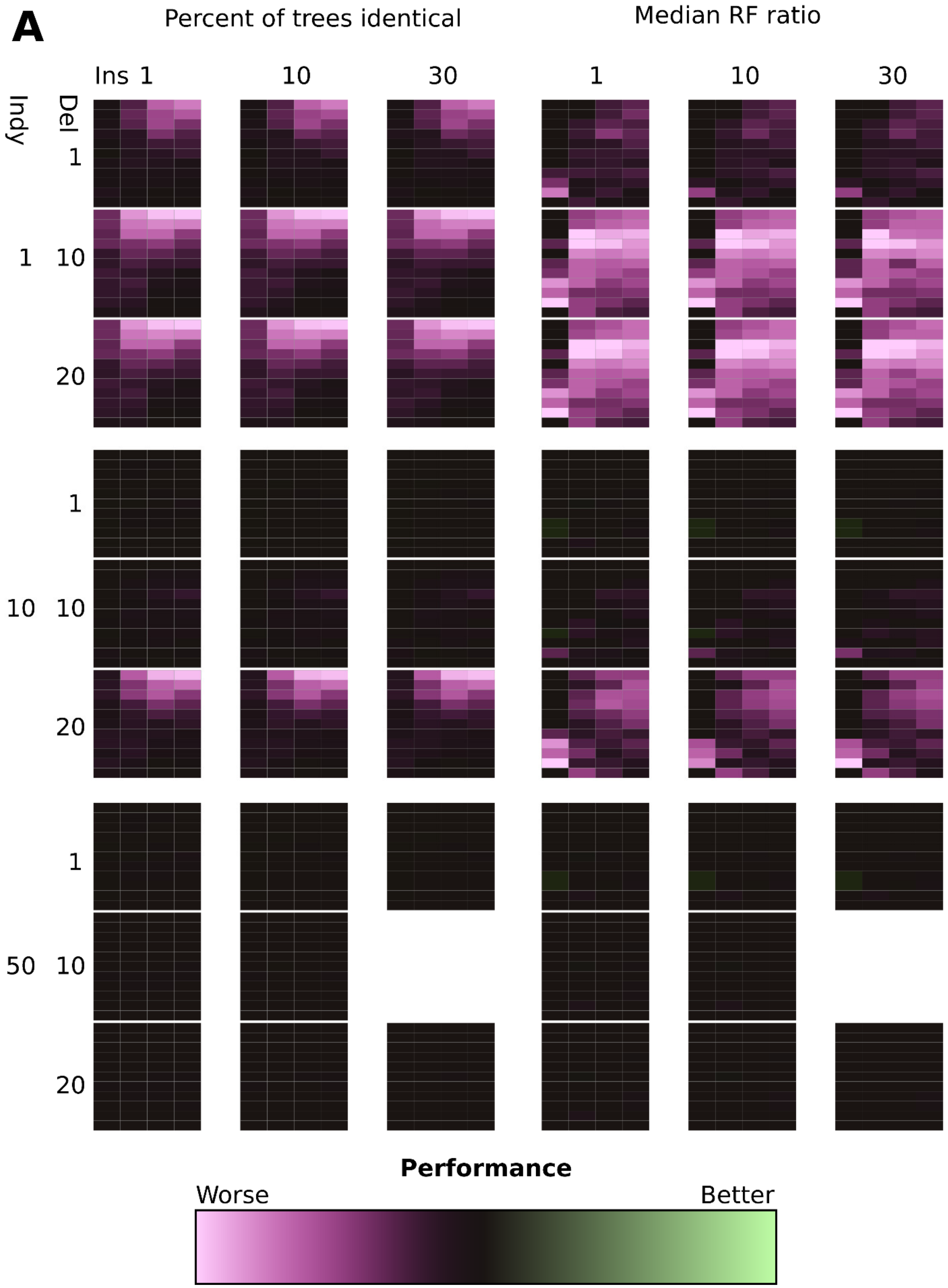
A
Percent of trees identical
Median RF ratio
Ins 1
10
30
1
10
30
Indy
Del
1
10
20
1
10
20
1
10
20
10
50
Performance
Worse
Better

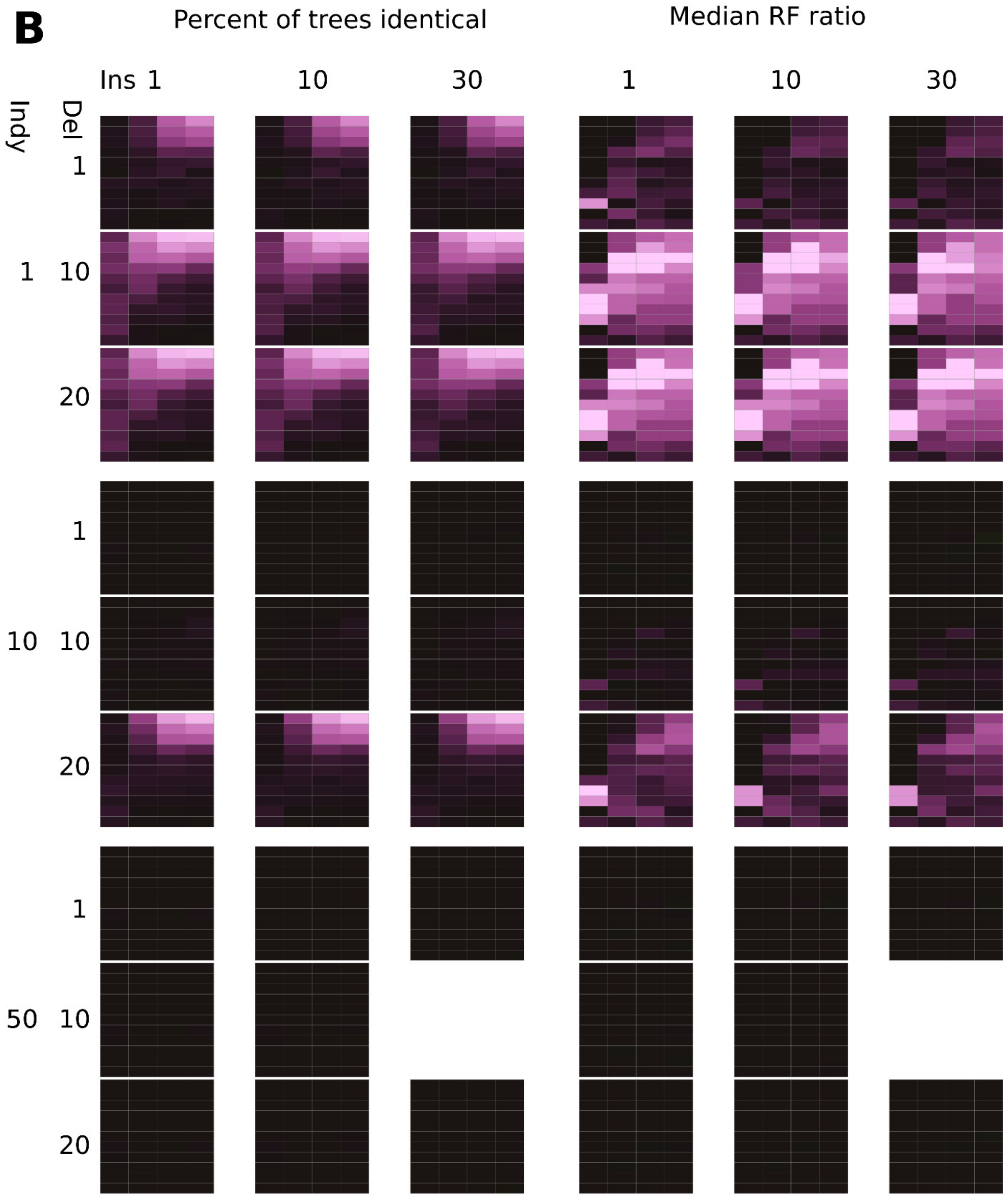
B
Percent of trees identical
Median RF ratio
Ins 1
10
30
1
10
30
Indy
Del
1
1
10
20
1
10
10
20
1
50
10
20

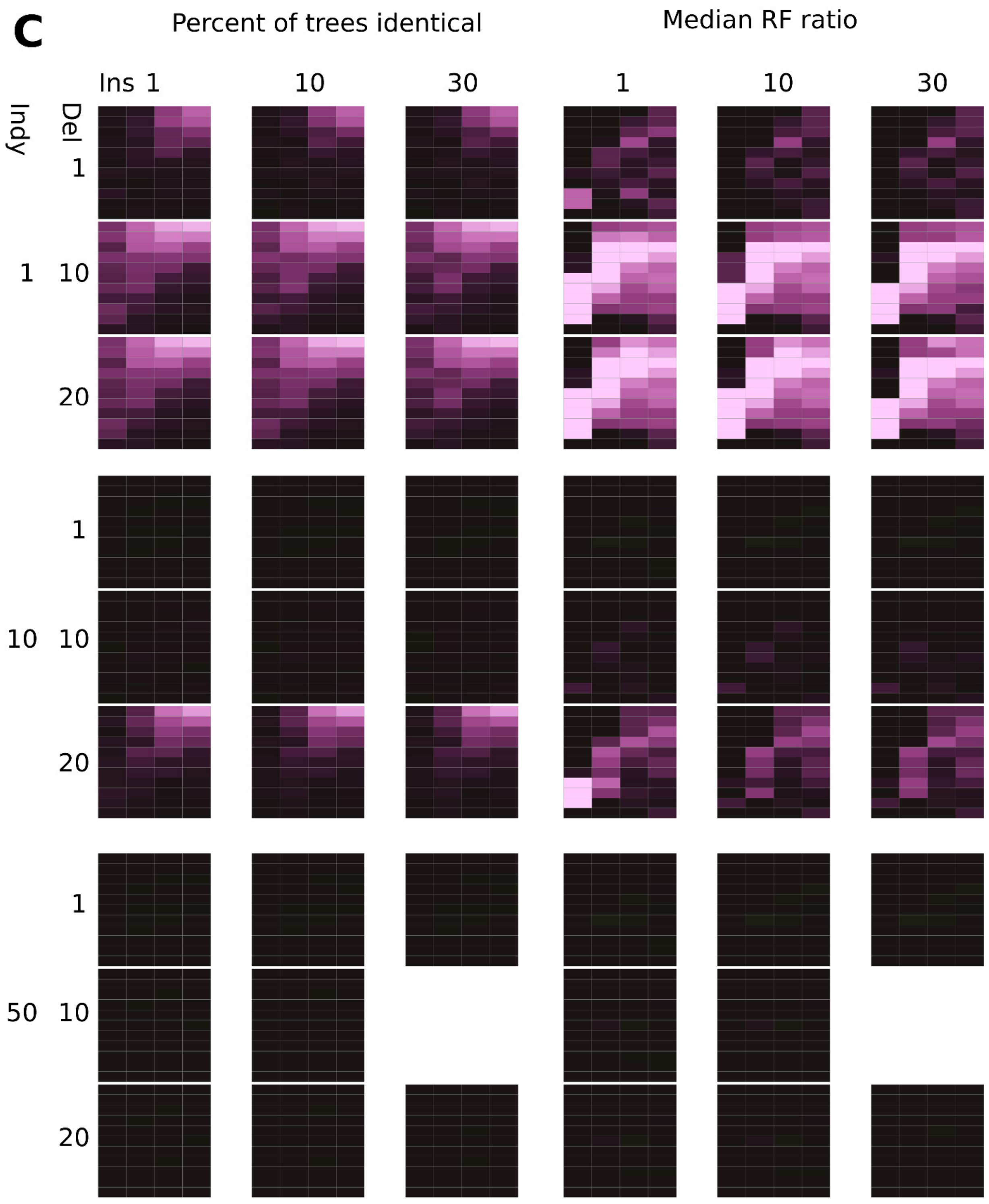
C
Percent of trees identical
Median RF ratio
Ins 1
10
30
1
10
30
Indy
Del
1
10
20
1
10
20
1
10
20
1
10
50

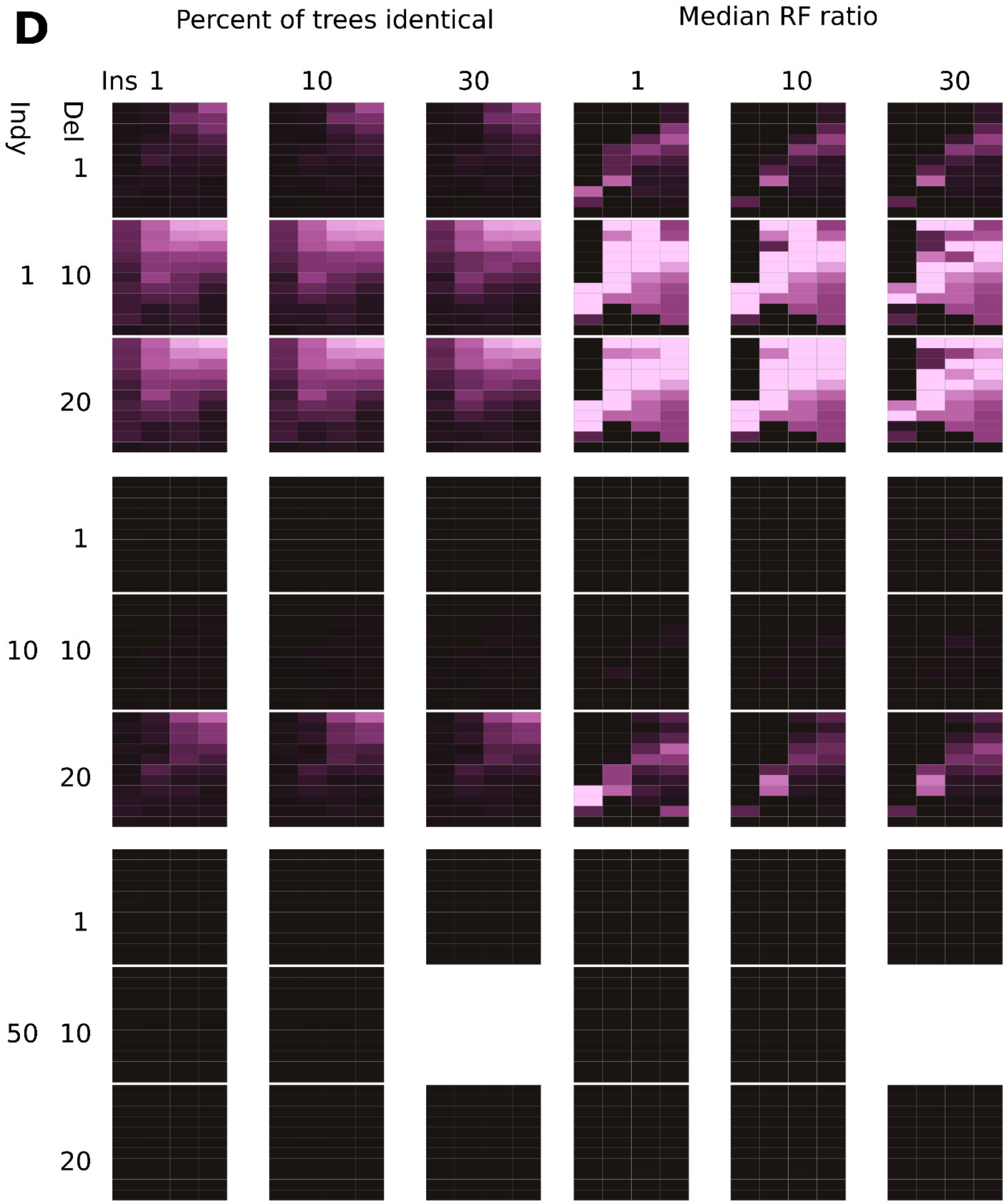
D
Percent of trees identical
Median RF ratio
Ins 1
10
30
1
10
30
Indy
Del
1
1 10
20
1
10 10
20
1
50 10
20

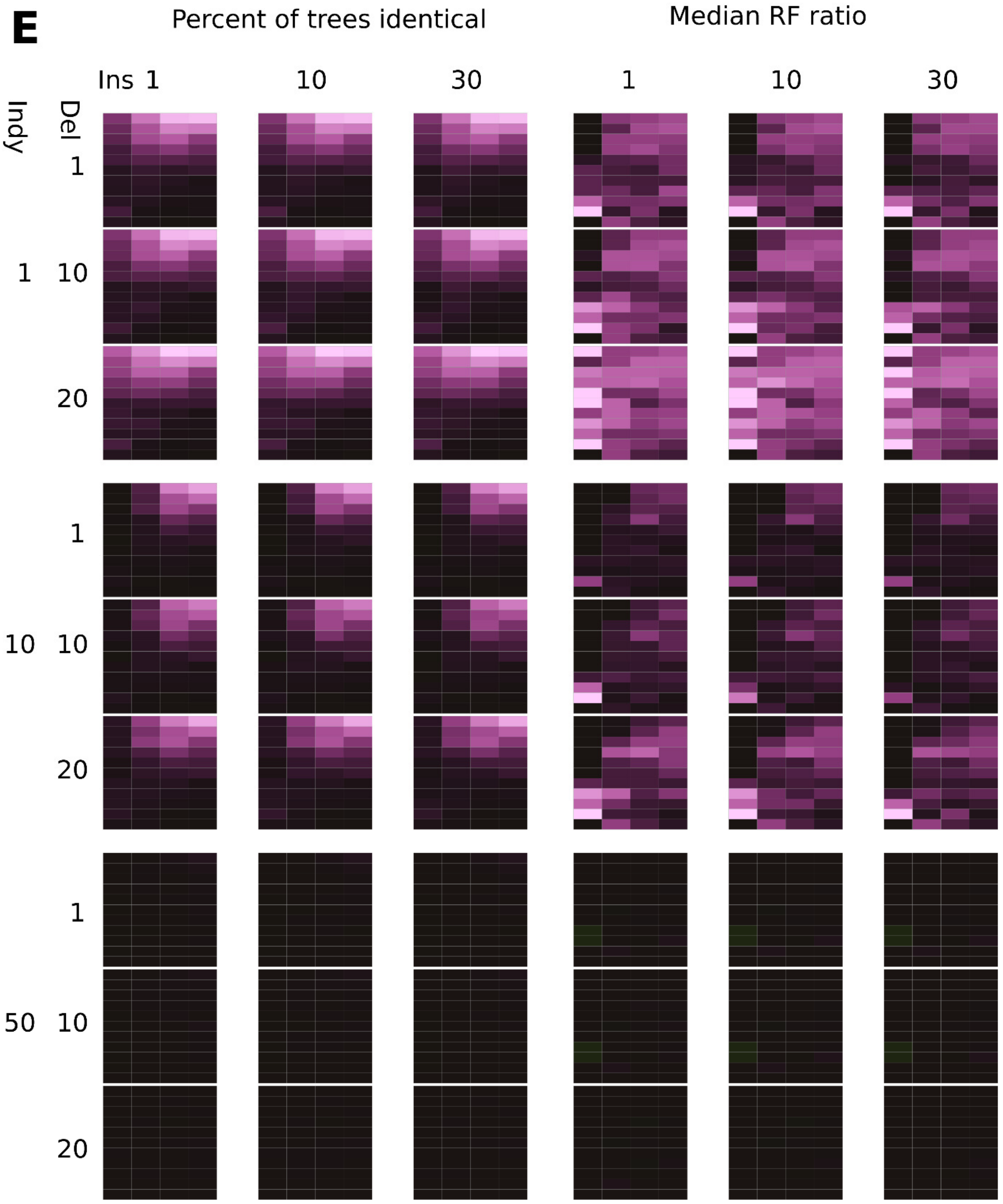
E
Percent of trees identical
Median RF ratio
Ins 1
10
30
1
10
30
Indy
Del
1
1
10
20
1
10
10
20
1
50
10
20

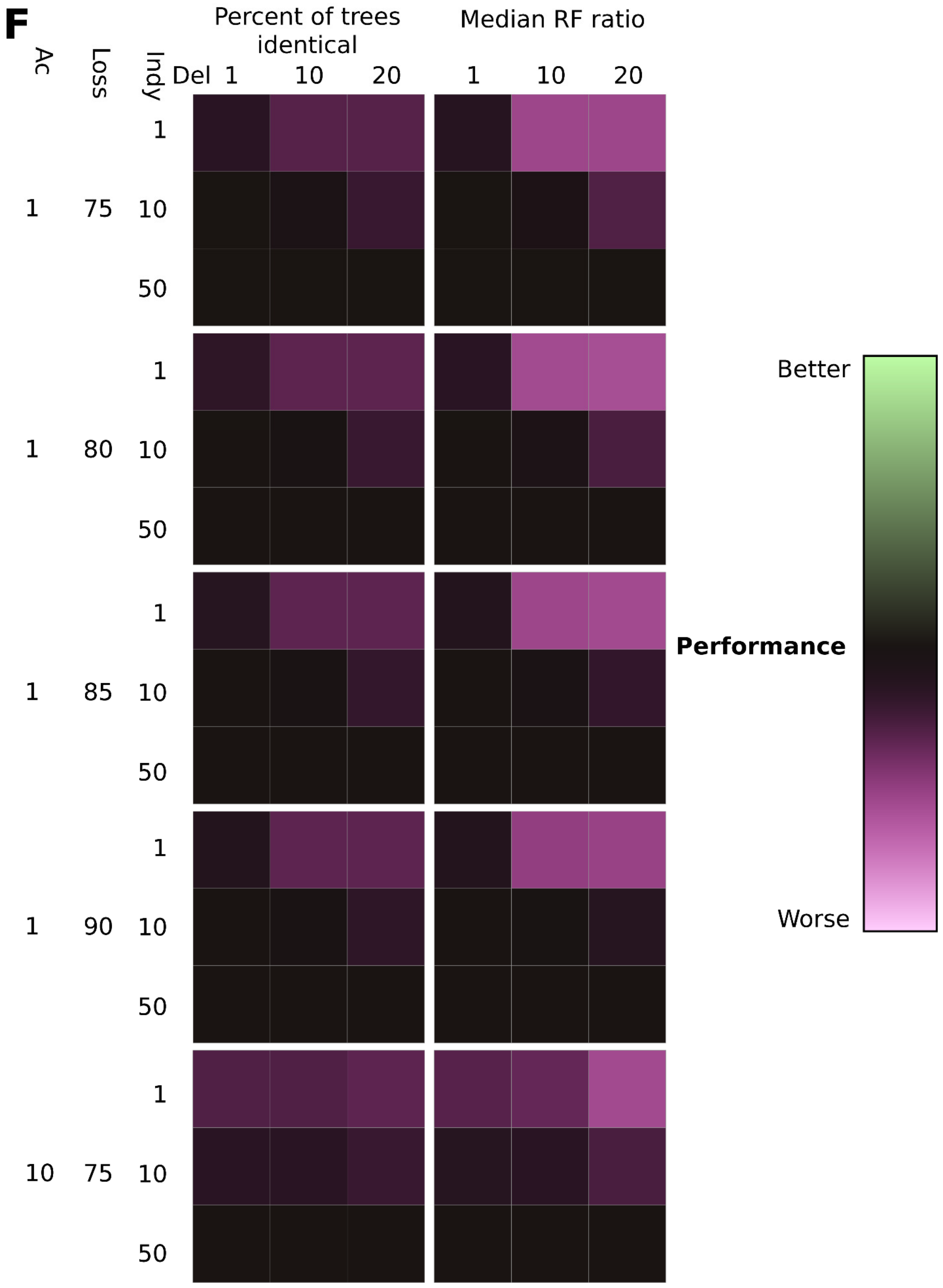
F
Percent of trees identical
Median RF ratio
Ac
Loss
Indy
Del
1
10
20
1
10
20
1
75
10
50
1
80
10
50
1
85
10
50
1
90
10
50
10
75
10
50
Better
Performance
Worse

**Figure S5. Impact of event parsimony cost on accuracy of CRISPRtree in reconstructing true tree topology.** (**A-D**) Relative performance of CRISPRtree assessed for simulated data with different loss rates: (**A**) 75%, (**B**) 80%, (**C**) 85%, (**D**) 90%. (**E**) Relative performance of CRISPRtree using an acquisition parsimony cost of 10 when analyzing data generated with a loss rate of 75% (compare to panel **A** in which loss rate is the 75% and acquisition cost is 1). In each panel (**A-D**), the heatmap corresponding to default parameters is omitted. (**F**) The mean value of each heatmap shown in panels **A-E** is summarized as a single value. Shown is the impact of the parsimony cost of deletions (columns of each heatmap) and independent acquisitions (rows of each heatmap) on CRISPRtree performance. Only heatmaps for insertion cost of 30 were summarized here.
